# Fungicide resistance characterised across seven modes of action in *Botrytis cinerea* isolated from Australian vineyards

**DOI:** 10.1101/2021.03.31.437981

**Authors:** Lincoln A. Harper, Scott Paton, Barbara Hall, Suzanne McKay, Richard P. Oliver, Francisco J. Lopez-Ruiz

## Abstract

**BACKGROUND:** Botrytis bunch rot, caused by *Botrytis cinerea*, is an economically important disease of grapes in Australia and across grape growing regions worldwide. Control of this disease relies on canopy management and the application of fungicides. Fungicide application can lead to the selection of resistant *B. cinerea* populations, which has an adverse effect on management of the disease. Characterising the distribution and severity of resistant *B. cinerea* populations is needed to inform resistance management strategies.

**RESULTS:** In this study, 725 isolates were sampled from 75 Australian vineyards during 2013 – 2016 and were screened against seven fungicides with different modes of action (MOAs). The resistance frequencies for azoxystrobin, boscalid, fenhexamid, fludioxonil, iprodione, pyrimethanil and tebuconazole were 5, 2.8, 2.1, 6.2, 11.6, 7.7 and 2.9% respectively. Nearly half of the resistant isolates (43.8%) were resistant to more than one of the fungicides tested. The frequency of vineyards with at least one isolate simultaneously resistant to 1, 2, 3, 4 or 5 fungicides was 19.5, 7.8, 6.5, 10.4 and 2.6%. Resistance was associated with previously published genotypes in *CytB* (G143A), *SdhB* (H272R/Y), *Erg27* (F412S), *Mrr1* (D354Y), *Bos1* (I365S, N373S + Q369P, I365S + D757N) and *Pos5* (V273I, P319A, L412F/V). Novel genotypes were also described in *Mrr1* (S611N, D616G) *Pos5* (V273L) and *Cyp51* (P347S). Expression analysis was used to characterise fludioxonil resistant isolates exhibiting overexpression (6.3-9.6-fold) of the ABC transporter gene *AtrB* (MDR1 phenotype).

**CONCLUSION:** Resistance frequencies were lower when compared to most previously published surveys of *B. cinerea* resistance in grape and other crops. Nevertheless, continued monitoring of critical MOAs used in Australian vineyards is recommended.

## 1 Introduction

*Botrytis cinerea Pers.:Fr.,* anamorph *Botryotinia fuckeliana (De Bary) Whetzel*, is a necrotrophic fungal pathogen with a broad host range. *B. cinerea* has been stated as only second to *Magnaporthe oryzae* (rice blast disease) in terms of its scientific and economic importance.^1^ There is currently a lack of scientific literature on the scale of crop losses caused by *B. cinerea*.^2^ *B. cinerea* is responsible for one of the most economically important diseases of grapevines worldwide. In Australia, *B. cinerea* is considered to be second only to powdery mildew in economic impact in grapes.^3^ *B. cinerea* infections, and to a lesser extent other bunch rots, impact all Australian grape growing regions and cost the grape and wine industry an average of $AUD50 M per annum.^3, 4^ Yield losses in Australia from *B. cinerea* and other bunch rots can be anywhere between 3 – 30 % depending on the climatic zone.^3^ *Botrytis* infection can also affect grape quality.^5^ The control of *B. cinerea* in vineyards relies heavily on the application of fungicides.^6^ In Australia, a wide range of both multi-site and single-site fungicides are registered for *B. cinerea* control. (Australian Pesticides and Veterinary Medicines Authority).

*B. cinerea* is a “high risk” pathogen for fungicide resistance development due to its short life cycle and high reproductive rate.^7, 8^ Resistance in *B. cinerea* has been linked to target site modifications, target site overexpression, efflux pump activation and detoxification. Resistance to single site fungicides was first reported in Germany in the 1970s after heavy use of dicarboximides (DCs) and benzimidazoles.^9^ To date, populations of *B. cinerea* from grapes resistant to the single site MOA anilinopyrimidines (APs), DCs, hydroxyanilides, phenylpyrroles (PPs), succinate dehydrogenase inhibitors (SDHIs and quinone outside inhibitor (QoI) classes, have been found in most grape growing countries.^10–20^ Resistance to these fungicides has been linked to target site modifications and efflux pump activation. In Australia, *B. cinerea* grapevine isolates resistant to benzimidazoles, DCs and APs have been previously described.^21, 22^ Similarly, a preliminary report showed isolates of *B. cinerea* from Australian vegetable crops have shown variable levels of resistance to SDHI, QoI and PP groups.^23^

The multiple drug resistance (MDR) phenotype is characterised by reduced sensitivity to fungicides and test compounds with diverse MOAs. MDR has been found in *B. cinerea* populations from grapevines, vegetables, and soft fruit crops.^24–28^ Two mechanisms that cause MDR in *B. cinerea* have been characterised; overexpression of the ABC transporter gene *AtrB* (MDR1 phenotype) and overexpression of the Major Facilitator Superfamily (MFS) transporter gene *MfsM2* (MDR2 phenotype).^29^

Current resistance management strategies for *Botrytis* include limiting fungicide use at the multi-seasonal scale, using mixtures, alternating MOA and the introduction of novel MOAs.^30^ For example, for APs and SDHIs, the Fungicide Resistance Action Committee (FRAC) recommends a maximum of three sprays per season. In Australia, a maximum of two single site fungicide sprays are recommended per season (https://www.awri.com.au/industry_support/viticulture/agrochemicals/agrochemical_booklet/). In Australia, MRLs (maximum residue levels) restrict the use of many fungicides on grapes used for the production of wine export.

The aim of this research was to provide data on fungicide sensitivity levels in *B. cinerea* Australian populations to seven fungicides classes widely used in wine grape production. A nationwide collection of isolates was screened via a combination of high-throughput discriminatory concentration assays and molecular analyses. In addition, the presence of MDR1 was investigated in selected isolates.

## 2 Materials and Methods

### 2.1 Fungal Isolates

During 2013-2016, mono-conidial isolates of *B. cinerea* were derived from infected grape material collected from 74 wine grape and 1 table grape vineyard across Australia (Table S1, Fig. 1 and 3). Sampling covered wine regions in Western Australia (WA), South Australia (SA), Victoria (VIC), New South Wales (NSW), Queensland (QLD) and Tasmania (TAS) (Table S1, Fig. 1). *B. cinerea* was sampled by directly swabbing infected material *in situ* or harvested material in the laboratory, with the swabs then used to inoculate yeast soluble starch medium, with addition of 1.25% w/v agar (YSSA).^31^ Isolates were subsequently single spored by using a sterile 25G hypodermic needle to transfer conidia to YSSA and were maintained as mycelial plugs on YSSA at 4 °C.

**Figure 1.**
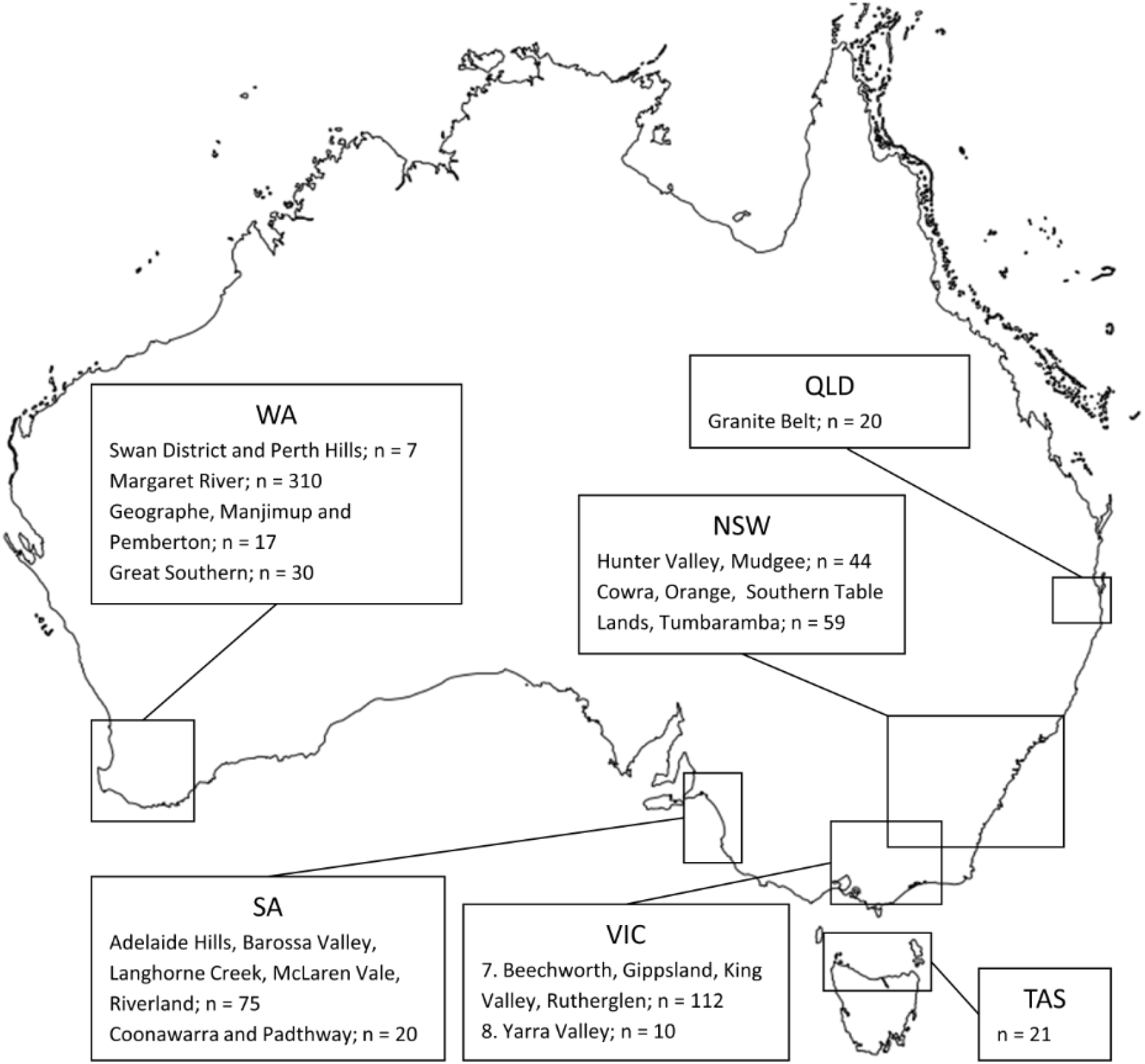
Distribution of the screened population (n = 725) across various Australian wine growing regions. Modified from “Outline map of Australia” (https://ecat.ga.gov.au/geonetwork/srv/eng/catalog.search#/metadata/61754) by Geoscience Australia, Canberra, used under CC BY 4.0.

### 2.2 Establishing baseline EC_50_ values for azoxystrobin, boscalid, fenhexamid, iprodione, pyrimethanil and tebuconazole using a microtiter assay

To establish baseline EC_50_ values, 53 isolates of *B. cinerea* sampled during 2013 – 2015 seasons, were randomly selected for testing against technical grade azoxystrobin (Sigma-Aldrich®, U.S.A), boscalid (Sigma-Aldrich®, U.S.A), fenhexamid (Bayer, Germany), iprodione (Sigma-Aldrich®, U.S.A), pyrimethanil (Bayer, Germany) and tebuconazole (Bayer, Germany) within a microtiter assay system (Table S2). Five isolates (Bc-7, Bc-279, Bc-287, Bc-385 and Bc-410) from the subset of 53 isolates, were selected for microtiter testing against fludioxonil (Sigma-Aldrich®, U.S.A). Additional isolates (Bc-247, Bc-296, Bc-298, Bc-398, Bc-403, Bc-477, Bc-618) were also tested against pyrimethanil using the microtiter method.

Induction of sporulation and the harvesting of conidia was carried out as previously described by Harper *et al*.^32^ with two modifications. YSS agar (YSSA) was used for culturing instead of potato dextrose agar and the conidial suspension adjusted to 10^5^ conidia mL^-1^, instead of 10^7^ conidia mL^-1^. The liquid media mixtures used in the microtiter assays were as described by Mair *et al*.,^33^ except with the addition of Tween® 20 at a final concentration of 0.05%, and the type of media used depended on the fungicide tested. Sensitivity to azoxystrobin, fenhexamid, fludioxonil, iprodione and tebuconazole was assessed using YSS medium, with the following fungicide concentrations: for azoxystrobin: 0, 0.01, 0.025, 0.05, 0.1, 0.5, 1 and 5µg mL^-1^, for fenhexamid: 0, 0.025, 0.05, 0.075, 0.1, 0.15, 0.25 and 1 µg mL^-1^, for iprodione: 0, 0.4, 1, 1.5, 2, 3, 5, and 10 µg mL^-1^, for fludioxonil: 0, 0.05, 0.1, 0.3, 0.6, 1, 1.5, 2, 2.5, 3, 5 and 10 µg mL^-1^, and for tebuconazole: 0, 0.2, 0.3, 0.5, 0.75, 1, 2, and 3 µg mL^-1^. Salicylhydroxamic acid (SHAM) (Sigma-Aldrich®, U.S.A) was also included in the media for azoxystrobin testing at a final concentration of 400 µM. Sensitivity to pyrimethanil was assessed using the amended YSS medium minus the yeast extract, with the following fungicide concentrations: 0, 0.05, 0.1, 0.2, 0.25, 0.3, 0.4 and 1 µg mL^-1^. Sensitivity to boscalid was assessed using Yeast Bacto Acetate (YBA) medium ^34^ with the following fungicide concentrations: 0, 0.01, 0.02, 0.03, 0.04, 0.075, and 0.1 µg mL^-1^. Re-testing of isolates that exhibited a significant reduction in sensitivity was carried out with at least one or more ranges of increased concentrations of fungicides; for azoxystrobin: 0, 0.5, 1, 5, 15, 25, 35 and 50 µg mL^-1^ or 0, 1, 5, 25, 50, 75, 100 and 150 µg mL^-1^, for boscalid: 0, 0.5, 1, 2, 3, 5, 7 and 10 µg mL^-1^, for fenhexamid: 0, 5, 10, 20, 30, 50, 75 and 100 µg mL^-1^, for iprodione: 0, 0.5, 1, 2, 3, 5, 7 and 10 µg mL^-1^ or 0, 1, 2, 3, 5, 6, 8 and 10 µg mL^-1^ or 0, 2, 4, 5, 7, 10, 25 and 50 µg mL^-1^, for : 0, 0.2, 0.5, 1, 1.5, 3, 5 and 10 µg mL^-1^ or 0, 0.2, 0.5, 1, 3, 5, 10 and 25 µg mL^-1^, for tebuconazole: 0, 0.25, 0.5, 1, 2, 3, 4 and 5 µg mL^-1^. The loading of conidia and media, reading of the microtiter plate, and the calculations of EC_50_ values was carried out as previously described by Mair *et al*.^33^, except that all plates were incubated for 72 h before reading at 450 nm with the exception of pyrimethanil plates that were read after 96h. EC_50_ values were calculated by linear regression of log_10_-transformed percentage inhibitions and fungicide concentrations. Resistance factors (RF) were calculated by dividing the resistant EC_50_ value by the mean EC_50_ value of the sensitive isolates.

### 2.3 Amplification and sequencing of the target genes; CytB, SdhB, Erg27, Bos1, Mdl1, Pos5, Cyp51, and the MDR1-related transcription factor Mrr1

To investigate mutations involved in resistance identified in the microtiter screen assays, all isolates that exhibited a significant reduction in sensitivity were genotyped for their respective resistance associated genes. The target genes for azoxystrobin, boscalid, fenhexamid, iprodione, pyrimethanil and tebuconazole were cytochrome B (*CytB*); succinate dehydrogenase subunit B (*SdhB*) and succinate dehydrogenase subunit B (*SdhD)*, 3-keto reductase (*Erg27*); histidine kinase (*Bos1*); mitochondrial ABC transporter (*Mdl1*) and mitochondrial NADH kinase (*Pos5*), and lanosterol 14 alpha-demethylase (*Cyp51*), respectively. Three sensitive strains (Bc-7, Bc-385 and Bc-410; Table S2) were genotyped for all target genes for comparative purposes. Candidate MDR1 isolates (Bc-128, Bc- 130, Bc-279, Bc-391) and the comparative isolate Bc-385, were sequenced for the *Mrr1* gene. The promoter for *Cyp51* was also sequenced for isolates exhibiting a reduction in sensitivity to tebuconazole. Additional isolates tested against pyrimethanil using the microtiter method (Bc-247, Bc-296, Bc-298, Bc-398, Bc-403, Bc-477, Bc-618), were genotyped for the *Pos5* gene. DNA was extracted from isolates as previously described by Harper *et al*.^32^ The primers used to amplify promoter and gene regions, and their respective annealing temperatures and extension times are shown in Table S3. All PCR reactions were carried out in an Eppendorf thermocycler model 5344. The *CytB* gene was amplified in a 50 µL reaction containing 2 µL of DNA, 10 µL of 5x HF reaction buffer, 0.16 mM of each dNTP, 0.5 µM of each primer and 1U of Phusion® polymerase (New England Biolabs® Inc., U.S.A). The subsequent thermal profile was as follows: initial denaturation was at 98 °C for 30 s followed by 35 cycles at 98 °C for 10 s, 59 °C for 30 s, and 72 °C for 3 min, and a final extension at 72 °C for 3 min. *SdhB, Erg27, Mrr1, Bos1, Mdl1, Pos5, Cyp51,* and the *Cyp51* promoter, were amplified using a MyTaq™ (Bioline, U.K.) reaction mixture and thermal profile as previously described by Harper *et al*^32^, except with the specific annealing temperatures and extension times as described in Table S3. Amplified gene products were confirmed on a 1% agarose gel and then sent to Macrogen Korea for sequencing. Consensus gene sequences were aligned to the following reference sequences: AB262969 (*CytB*), AY726618 (*SdhB*), GQ253439 (*SdhD*), AY220532 (*Erg27*), *B. cinerea* B05.10 chromosome 5; CP009809 (*Mrr1*), AF435964 (*Bos1*), *B. cinerea* B05.10 chromosome 10; CP009814 (*Pos5*), *B. cinerea* B05.10 chromosome 16; CP009820 (*Mdl1*), AF279912 (*Cyp51*), *B. cinerea* T4 contig; FQ790352 (*Cyp51* promoter). Alignments were carried out as described by Mair *et al*.^33^. The nucleotide sequences generated in this study have been deposited in GenBank and are listed in Table S4.

### 2.4 Cleaved amplified polymorphic sequence analysis of Bos1

The cleaved amplified polymorphic sequence (CAPS) test utilising the restriction enzyme *Taq* I (New England Biolabs® Inc., U.S.A) to identify the I365N/R/S mutant in *Bos1* was carried out as described by Oshima *et al*.^35^ In this study, the internal sequencing primers os1 F and os1 R (Table S3) were used to amplify a 1133 bp fragment encompassing the I365S and Q369P + N373S alleles. The fragment was amplified using the MyTaq protocol described above and with the annealing temperature and extension time described in Table S3. The digestion of the product was carried as per Oshima *et al*.^35^ The same amplified fragment was also used to test for the presence of the Q369H/P allele, which required digestion with *Sma* I (New England Biolabs® Inc., U.S.A).

### 2.5 Development of a fungicide resistance discriminatory concentration agar assay

Minimum inhibitory concentration (MIC) values identified in the microtiter assay (Table S2) were used to design a discriminatory concentration (DC) agar screen to test the remaining 672 isolates in the collection. The DCs used in the agar assay for azoxystrobin, boscalid, fenhexamid, fludioxonil, iprodione, pyrimethanil and tebuconazole were 5, 1, 1, 0.1, 3, 0.4 and 3 µg mL^-1^. For all fungicides, the DC screening was carried out using YSS agar, except for boscalid which used YBA agar and pyrimethanil which used YSSA minus the yeast extract. SHAM was added at a concentration of 400 µM for the plates containing azoxystrobin. Mycelial colonies were grown as described by Harper *et al*.^32^. Two 4 mm agar plugs per isolate were taken from the edge of actively growing colonies and then placed on plates containing the appropriate media and fungicide combination. Eight isolates were tested per plate, and plates were incubated at room temperature in the dark for 3 days and scored based on their ability to grow on each fungicide.

### 2.6 RT-qPCR analysis of AtrB and Cyp51

To identify MDR1 phenotypes and further characterise isolates lacking mutations in *Cyp51*, the expression levels of *AtrB* and *Cyp51* were assessed, respectively. Isolates Bc-128, Bc-130, Bc-279, Bc- 287, Bc-385 and Bc-391 were selected for *AtrB* expression analysis. Bc-130 was selected for *AtrB* analysis to investigate if MDR1 could contribute to a teb^LR^ phenotype. Bc-287 was selected for *AtrB* expression analysis as it exhibited a group S haplotype (Fig. 8) that has previously been associated with MDR1.^24–26^ Isolates were incubated as described by Li *et al*.,^25^ with some modifications. Culture growth was carried out in 50 mL of potato dextrose broth (Difco™, U.S.A) in a 250 mL flask. For *AtrB* expression analysis, Bc-279, Bc-287, and Bc-385 were induced with 1 µg mL^-1^ pyrimethanil for 0.5 h. For *Cyp51* expression analysis, Bc-130 and Bc-385 were induced with 0.5 µg mL^-1^ tebuconazole for 1 h. Primers that were used for RT-qPCR analysis are described in Table S3. Harvesting of biomass, extraction of RNA, production of cDNA, and RT-qPCR was carried out as described by Mair *et al*.^33^ with *Actin* used as the endogenous control (BC1G_08198.1).

### 2.7 Statistical analysis

All statistical analyses were conducted in SPSS statistics (IBM, New York, U.S.A.). To account for heteroscedasticity, all fungicide sensitivity data were log_10_-transformed before analysis. To separate EC_50_ mean values between sensitive and resistant isolates characterised in the 53 isolate microtiter subset, an independent samples t-test (*P* = 0.05) or Mann-Whitney *U*-test (*P* = 0.05) was used depending on whether the data set had a normal or non-normal distribution. Mean values for gene expression analyses were analysed using one-way ANOVA, with means separated by independent samples t-test (*P* = 0.05)

## 3 Results

### 3.1 Identification of resistance to multiple fungicide groups in B. cinerea

The EC_50_ ranges for azoxystrobin, boscalid, fenhexamid, fludioxonil, iprodione, pyrimethanil and tebuconazole, were 0.04–>50, 0.03–2.90, 0.05–27.33, 0.09 – 0.93, 0.75–>50, 0.09–28.71 and 0.21– 1.8 µg mL^-1^, respectively (Tables 1 and S2, Fig. 2). Except for iprodione and tebuconazole, isolates with an EC_50_ >0.5 µg mL^-1^ were classified as resistant. For iprodione and tebuconazole, isolates were considered resistant when their EC_50_ values were higher than 2.5 µg mL^-1^ and 0.75 µg mL^-1^, respectively (Tables 1 and S2, Fig. 2). Isolates resistant to azoxystrobin, boscalid and fenhexamid were all designated as azo^R^, bos^R^ and fen^R^ isolates, respectively (Tables 1 and S2, Fig. 2). Isolates resistant to iprodione, fludioxonil and pyrimethanil were divided into medium resistant (ipr^MR^, flu^MR^, pyr^MR^) or high resistant (ipr^HR^, pyr^HR^) isolates (Tables 1 and S2, Fig. 2). Tebuconazole resistant isolates were characterised as low resistant (teb^LR^) (Tables 1 and S2, Fig. 2). EC_50_ values for the sensitive and resistant populations for all fungicides were significantly difference from each other (*P* < 0.05; Tables 1 and S2, Fig. 2). The RF ranges of the resistant isolates in the 53 isolate subset ranged from low (2.5 – 4.2) for tebuconazole, medium (4.3 - 58) for boscalid, iprodione and pyrimethanil, and high (>255) for azoxystrobin and fenhexamid (Table 1).

**Figure 2.**
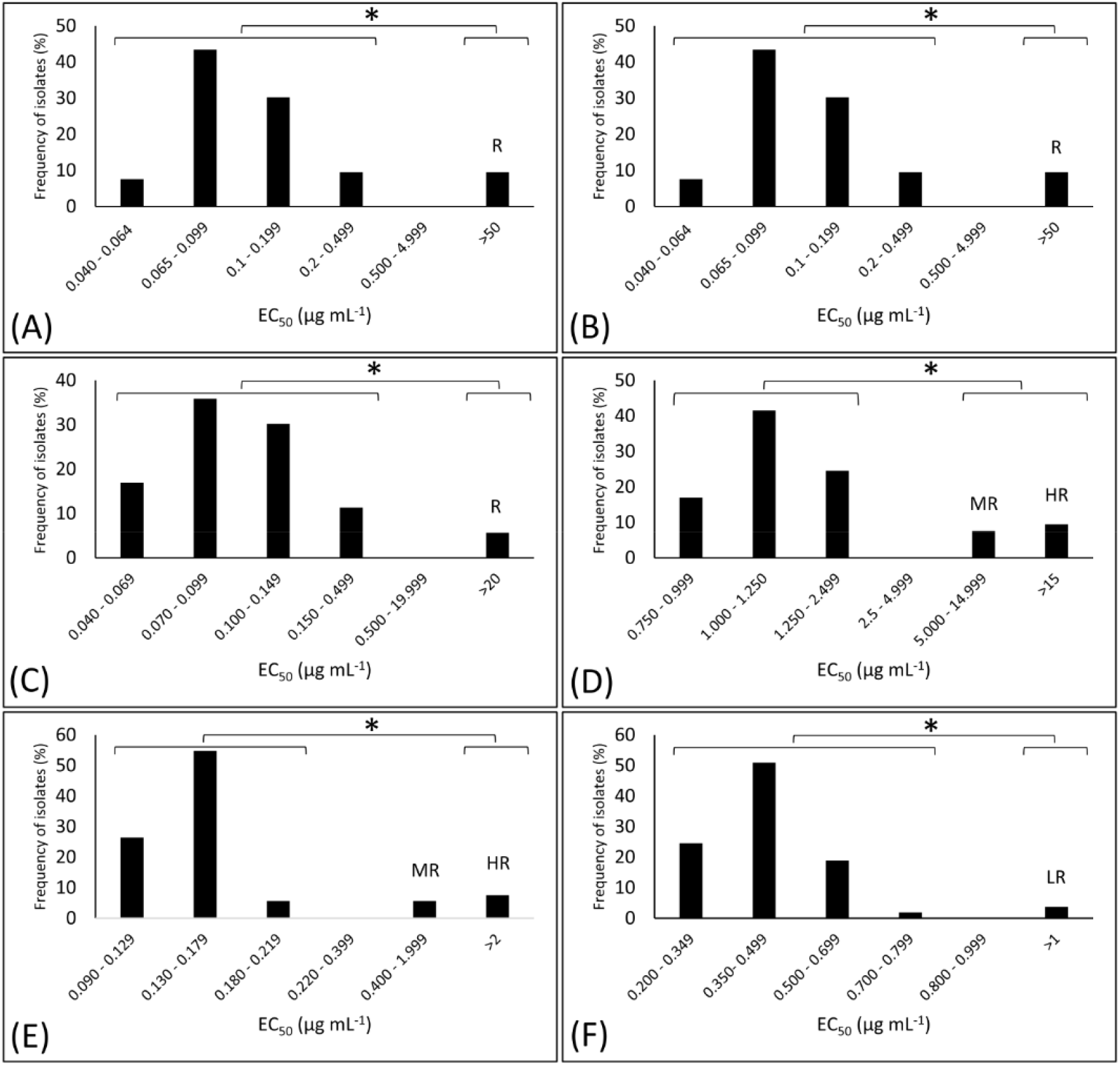
EC_50_ distribution of 53 isolates phenotyped in the microtiter assay against azoxystrobin (A), boscalid (B), fenhexamid (C), iprodione (D), pyrimethanil (E), and tebuconazole (F). LR, MR, HR and R indicate isolates designated as exhibiting low resistance, medium resistance, high resistance, or resistance, respectively. *indicates significant difference between the means of the sensitive and resistant populations according to an independent samples t-test (*P* = 0.05) or Mann-Whitney U-test (*P* = 0.05).

**Table 1.** Results summary of the microtiter phenotyping of 53 *B. cinerea* isolates.

|  |  | Azoxystrobin | Boscalid | Fenhexamid | Iprodione | Pyrimethanil | Tebuconazole |
| --- | --- | --- | --- | --- | --- | --- | --- |
| EC<sub>50</sub> (µg mL<sup>-1</sup>) | Sensitive range | 0.04 - 0.37 | 0.03 - 0.10 | 0.05 - 0.22 | 0.75 - 2.05 | 0.09 - 0.22 | 0.21 - 0.72 |
|  | Sensitive mean | 0.12 ± 0.010 | 0.05 ± 0.002 | 0.10 ± 0.006 | 1.20 ± 0.039 | 0.14 ± 0.004 | 0.43 ± 0.015 |
|  | Resistant range | >50 | 2.74 - 2.90 | 25.47 - 27.33 | 5.16 - >50 | 0.75 - 5.90 | 1.06 - 1.80 |
|  | Resistant mean<sup>a</sup> | 171.26 ± 16.264 | 2.82 ± 0.076 | 26.19 ± 0.578 | 21.80 ± 4.901 | 2.99 ± 0.805 | 1.43 ± 0.372 |
|  | LR range | - | - | - | - | - | 1.06 - 1.80 |
|  | MR range | - | - | - | 5.16 - 11.94 | 0.75 - 2.27 | - |
|  | HR or R range | >50 | 2.74 - 2.90 | 25.46 - 27.33 | >16.78 | 4.66 - 5.90 | - |
| no. of isolates | Sensitive | 48 | 51 | 50 | 43 | 46 | 51 |
|  | Resistant | 5 | 2 | 3 | 9 | 7 | 2 |
|  | RF range | >413 | 55 - 58 | 255 - 273 | 4.3 - >42 | 5 - 42 | 2.5 - 4.2 |
|  | Resistance frequency (%) | 9.3 | 3.7 | 5.7 | 17 | 13.2 | 3.8 |

<sup>a</sup>: Significant differences were found between the resistant and sensitive populations according to independent samples t-test ( $P = 0.05$ ) or Mann-Whitney U-test ( $P = 0.05$ ).

### 3.2 Resistance frequencies for seven MOAs in Australian grape growing regions

A total of 672 isolates collected between 2013 and 2016 from major grape-producing regions of Australia were screened for resistance using a DC agar assay as described above. Combining microtiter and DC agar assay data, the total resistance frequencies ranged from 2.8 to 11.6% (Fig. 3). Thirty different resistance profiles, with 24 of these showing resistance to at least 2 MOA, were identified (Table 2). The total number of resistant isolates for each fungicide in WA, SA, VIC and NSW, ranged from 0 – 30, 3 – 14, 1 – 26 and 0 – 3, respectively (Fig. 4). The frequency of resistant isolates in each of the states was 15.4, 25.3, 61.9, 36.9, and 5.8% across WA, SA, TAS, VIC and NSW, respectively (Table S1). The frequency of vineyards with resistance to 1 – 5 MOA were 19.5, 7.8, 6.5, 10.4 and 2.6%, respectively (Fig. 5). No resistance was found in QLD.

**Figure 3.**
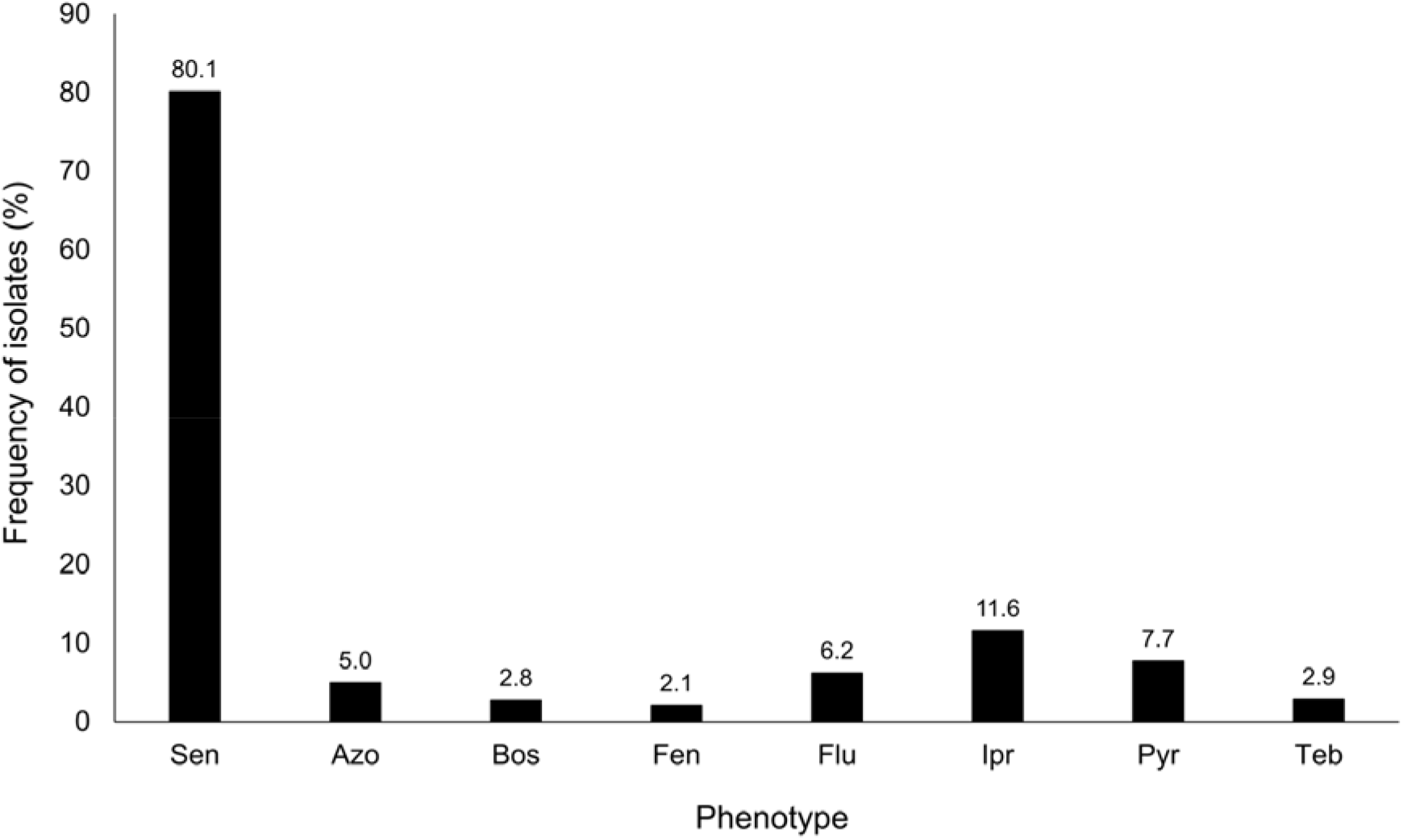
Frequency (%) of resistant isolates in the population (n=725) on a per fungicide basis. Azo = azoxystrobin resistance, Bos = boscalid resistance, Fen = fenhexamid resistance, Pyr = pyrimethanil resistance, Ipr = iprodione resistance, teb = tebuconazole resistance. Percentage for each fungicide is shown above each column. These results do not consider resistance to more than one MOA.

**Figure 4.**
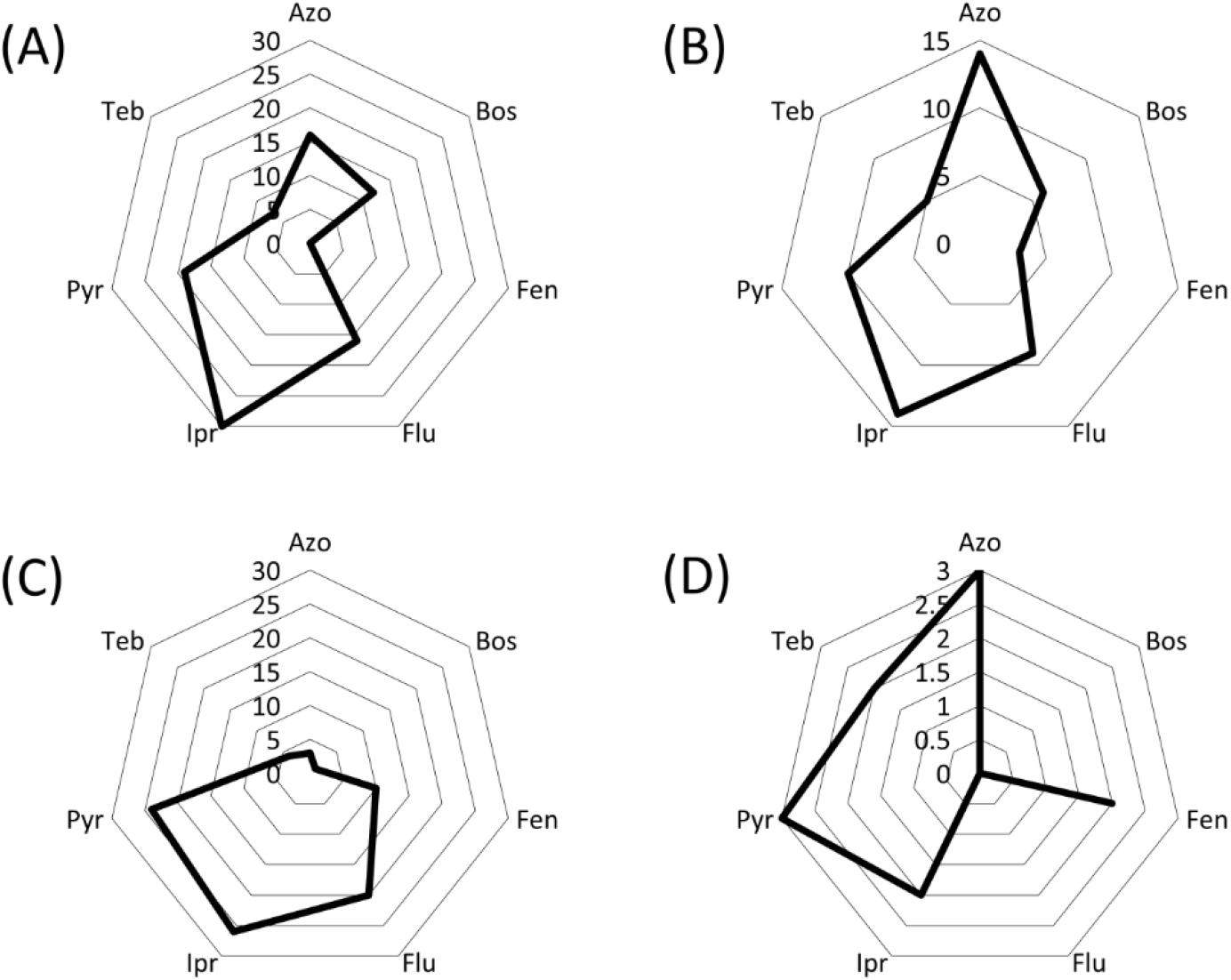
Number of resistant isolates in each state on a per fungicide basis. A = Western Australia, B = South Australia, C = Victoria, D = New South Wales. Tasmania and Queensland were omitted due to small sample size (≤ 21). Azo = azoxystrobin resistance, Bos = boscalid resistance, Fen = fenhexamid resistance, Pyr = pyrimethanil resistance, Ipr = iprodione resistance, Teb = tebuconazole resistance.

**Figure 5.**
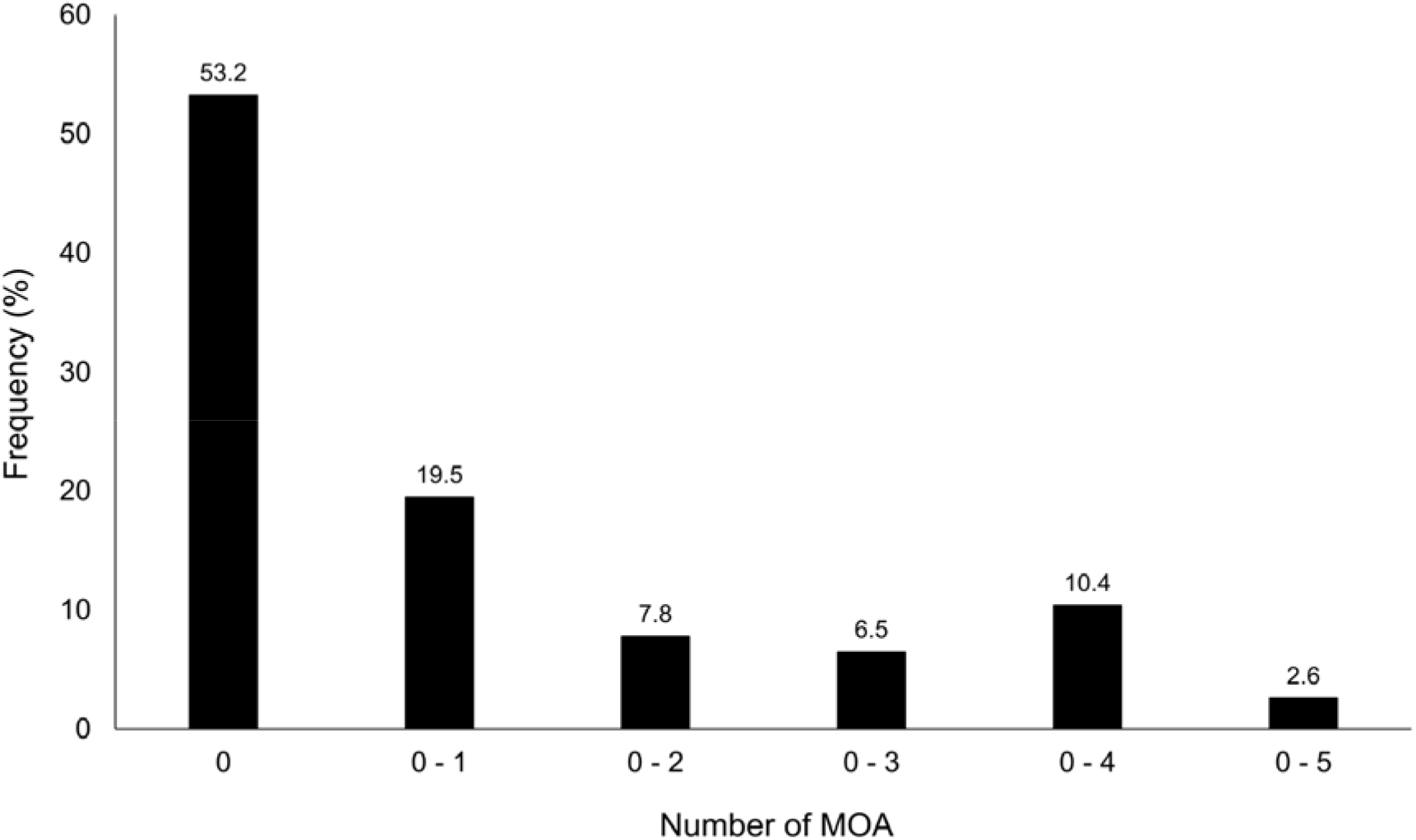
Frequency (%) of vineyards with no resistance and with at least one isolate resistant to one to five modes of action (MOAs). Percentage for each grouping is shown above each column.

**Table 2.** Frequency of phenotypes characterised across six Australian states.

|  | WA | SA | TAS | VIC | NSW | QLD | Total (%) |
| --- | --- | --- | --- | --- | --- | --- | --- |
| Isolates tested | 364 | 95 | 21 | 121 | 103 | 20 | 725 |
| Vineyards sampled | 38 | 13 | 1 | 11 | 13 | 1 | 77 |
| Total sensitive | 308 (84.6) | 71 (74.7) | 8 (38.2) | 76 (62.8) | 97 (94.2) | 20 (100) | 581 (80.1) |
| Resistance phenotype<sup>a</sup> |  |  |  |  |  |  |  |
| Azo | 1 | 1 | 0 | 0 | 1 | 0 | 3 (0.4) |
| Bos | 2 | 0 | 1 | 0 | 0 | 0 | 3 (0.4) |
| Flu | 11 | 1 | 0 | 10 | 0 | 0 | 22 (3) |
| Ipr | 14 | 2 | 9 | 9 | 1 | 0 | 35 (4.8) |
| Pyr | 1 | 0 | 0 | 5 | 1 | 0 | 7 (1) |
| Teb | 5 | 4 | 0 | 1 | 1 | 0 | 11 (1.5) |
| Azo Bos | 0 | 2 | 0 | 0 | 0 | 0 | 2 (0.3) |
| Azo Flu | 0 | 1 | 0 | 0 | 0 | 0 | 1 (0.1) |
| Azo Ipr | 2 | 0 | 0 | 0 | 0 | 0 | 2 (0.3) |
| Azo Pyr | 2 | 0 | 0 | 1 | 0 | 0 | 3 (0.4) |
| Flu Ipr | 1 | 1 | 0 | 1 | 0 | 0 | 3 (0.4) |
| Flu Pyr | 0 | 0 | 0 | 1 | 0 | 0 | 1 (0.1) |
| Flu Teb | 1 | 1 | 0 | 0 | 0 | 0 | 2 (0.3) |
| Ipr Pyr | 2 | 0 | 0 | 1 | 0 | 0 | 3 (0.4) |
| Ipr Teb | 0 | 0 | 3 | 0 | 0 | 0 | 3 (0.4) |
| Pyr Teb | 0 | 0 | 0 | 1 | 0 | 0 | 1 (0.1) |
| Azo Ipr Pyr | 1 | 1 | 0 | 0 | 0 | 0 | 2 (0.3) |
| Azo Bos Pyr | 3 | 0 | 0 | 0 | 0 | 0 | 3 (0.4) |
| Azo Fen Pyr | 0 | 0 | 0 | 0 | 1 | 0 | 1 (0.1) |
| Azo Flu Ipr | 0 | 1 | 0 | 0 | 0 | 0 | 1 (0.1) |
| Fen Ipr Pyr | 0 | 0 | 0 | 5 | 0 | 0 | 5 (0.7) |
| Flu Ipr Pyr | 2 | 0 | 0 | 2 | 0 | 0 | 4 (0.6) |
| Azo Bos Ipr Pyr | 7 | 2 | 0 | 1 | 0 | 0 | 10 (1.4) |
| Azo Fen Ipr Pyr | 0 | 2 | 0 | 1 | 0 | 0 | 3 (0.4) |
| Azo Flu Ipr Pyr | 0 | 3 | 0 | 0 | 0 | 0 | 3 (0.4) |
| Bos Flu Ipr Pyr | 0 | 1 | 0 | 0 | 0 | 0 | 1 (0.1) |
| Fen Flu Ipr Pyr | 0 | 0 | 0 | 4 | 0 | 0 | 4 (0.6) |
| Flu Ipr Pyr Teb | 1 | 0 | 0 | 2 | 0 | 0 | 3 (0.4) |
| Azo Bos Fen Ipr Pyr | 0 | 1 | 0 | 0 | 0 | 0 | 1 (0.1) |
| Azo Fen Ipr Pyr Teb | 0 | 0 | 0 | 0 | 1 | 0 | 1 (0.1) |
| Total resistant (%) | 56 (15.4) | 24 (25.3) | 13 (61.9) | 45 (37.2) | 6 (5.8) | 0 | 144 (19.8) |

<sup>a</sup>: Azo = azoxystrobin, Bos = boscalid, Fen = fenhexamid, Ipr = iprodione, Pyr = pyrimethanil, Teb = tebuconazole

### 3.3 Fungicide resistance is associated with mutations in multiple fungicide target genes

Relevant target site genes were sequenced in all resistant isolates found in the microtiter assay and compared to target gene sequences from the three sensitive reference isolates (Bc-7, Bc-385, Bc- 410) (Table S2, Fig. 6). Mutations in archetype species are stated below if the proposed archetype is not *B. cinerea* or if the mutations are not exclusively characterised in *B. cinerea*.^36^

**Figure 6.**
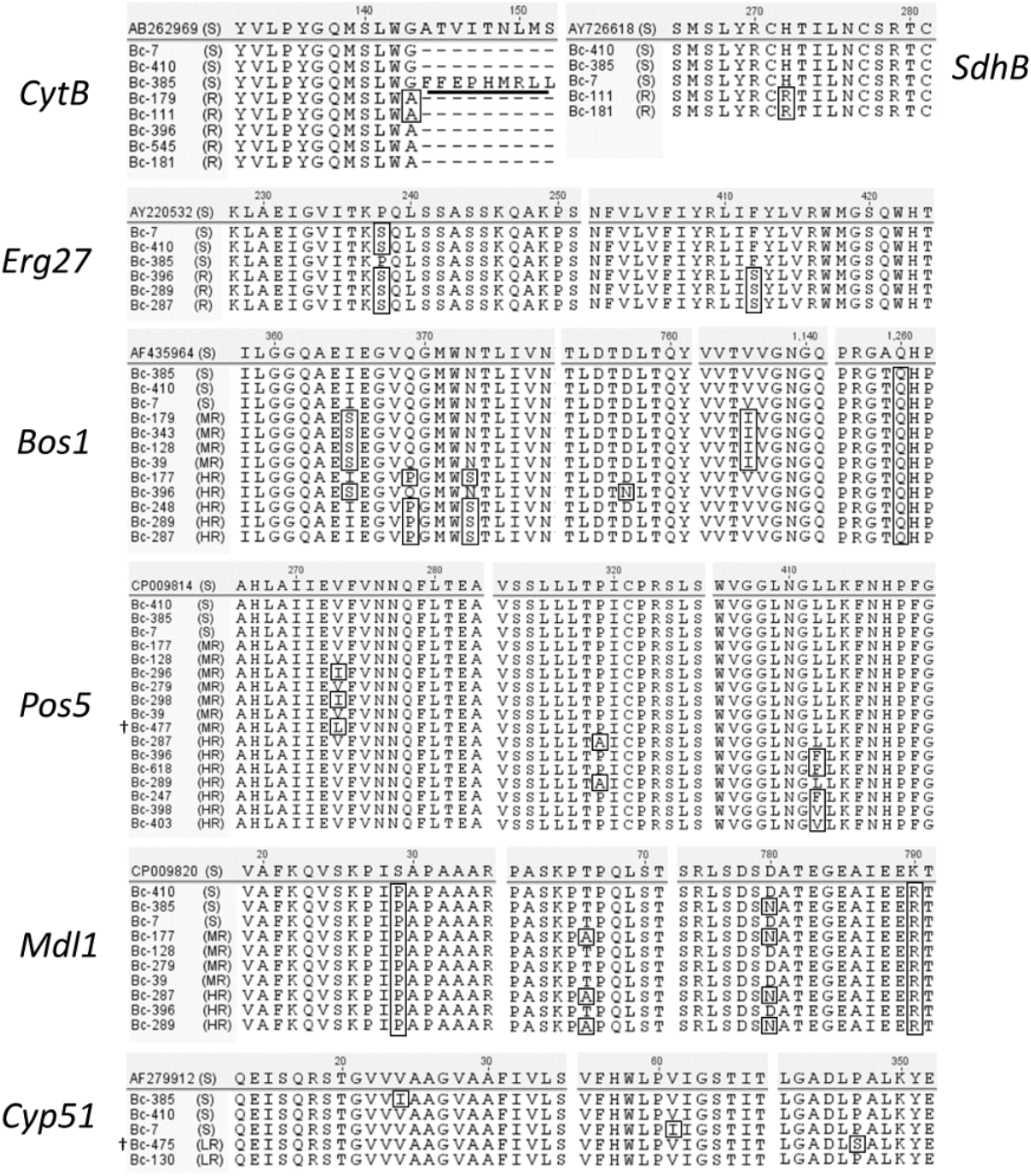
Amino acid sequence alignment of fungicide target genes; *CytB* (azoxystrobin), *SdhB* (boscalid), *Erg27* (fenhexamid), *Bos1* (iprodione), *Pos5* (pyrimethanil), *Mdl1* (pyrimethanil), *Cyp51* (tebuconazole), for three sensitive comparative isolates and all resistant isolates characterised in the 53 isolate microtiter assay screen. For each target gene, isolates are ordered from most sensitive to the least sensitive in a descending order. LR, MR, HR and R indicate isolates designated as exhibiting low resistance, medium resistance, high resistance, or resistance, respectively. Black boxes represent non-synonymous changes found when compared to the reference sequence. The *CytB* G143 intron is underlined in Bc-385. † = isolates with novel genotypes.

Azo^R^ isolates all showed the amino acid sequence change G143A (G143A in the archetype *Zymoseptoria tritici*) (Table 3, Fig. 6). Sequencing of *CytB* from the sensitive isolate Bc-385 revealed the presence of an intron at the G143 site (Fig. 6). Amplification of *CytB* from Bc-111 revealed the presence of two fragments, indicating the presence of the G143 intron (data not shown). Subsequently, intron specific primers (Table S3) were used to confirm the presence of this intron in Bc-111 (data not shown).

**Table 3.**
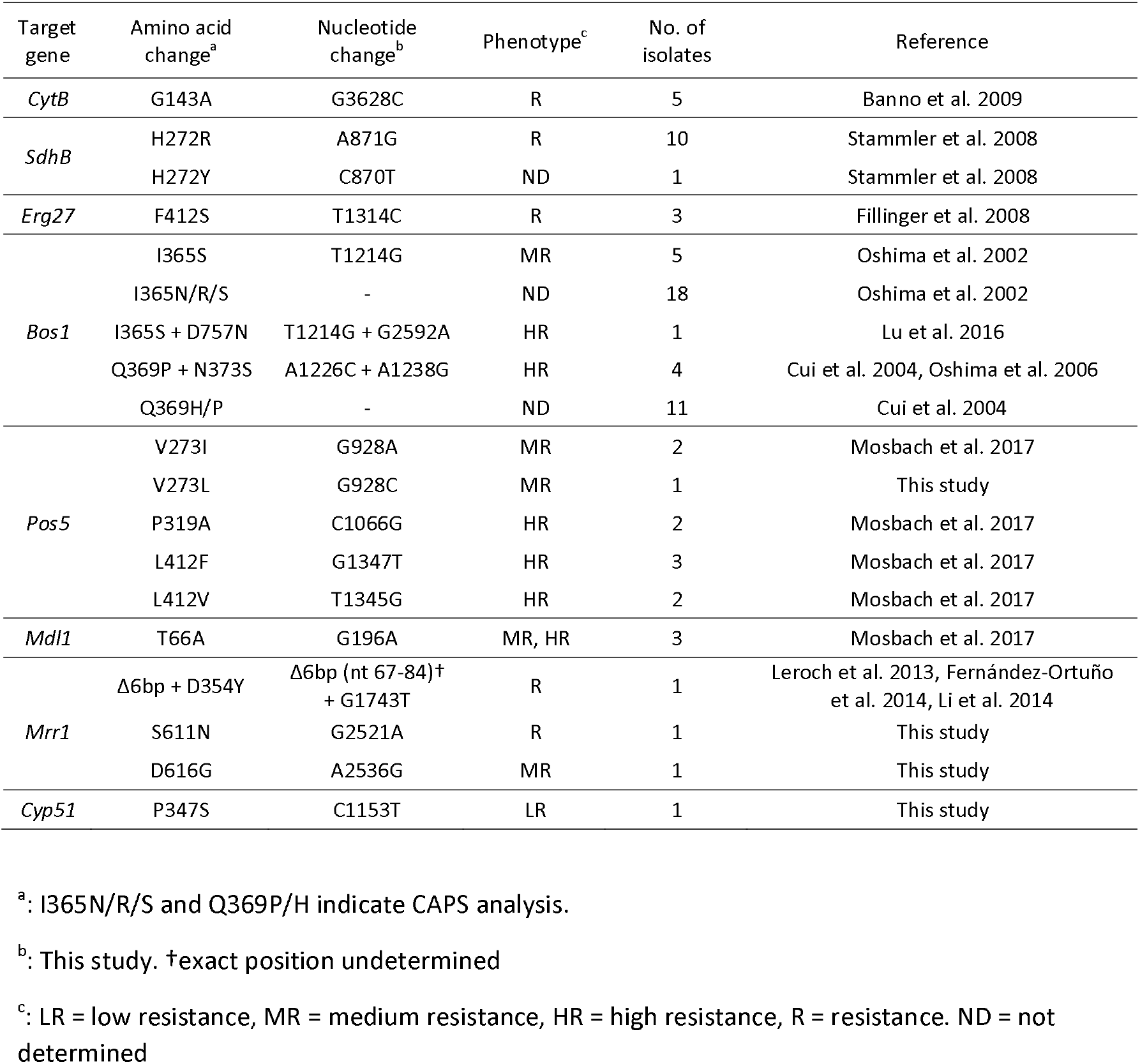
Frequency of mutations associated with resistance identified in this study.

Sequencing of the *SdhB* gene in two bos^R^ isolates identified in the microtiter analysis showed mutation H272R (*H277Y* in the archetype *Pyrenophora teres f. sp. teres*) (Table 3, Fig. 6). Additional *SdhB* genotyping was carried out on nine boscalid resistant isolates identified in the DC assay (Table 3). Eight of these isolates contained the H272R mutation, while the remaining isolate had the amino acid change H272Y (Table 3, Fig. 6). Sequencing of the *SdhD* gene in two bos^R^ isolates (Bc-111, Bc- 181) and comparative sensitive isolates revealed no non-synonymous changes (data not shown).

A polymorphism in *Erg27* at amino acid position 238 (CCT to TCT) that resulted in a change of a proline to a serine (P238S) was not correlated with fenhexamid resistance (Fig. 6). Fen^R^ isolates had the amino acid change F412S (Table 3, Fig. 6).

All sensitive and resistant isolates genotyped for *Bos1* had the amino acid change A1259T (Fig. 6). All ipr^MR^ isolates had the amino acid changes I365S and V1136I (Table 3, Fig. 6). All ipr^HR^ isolates either contained the amino acid changes Q369P + N373S or I365S + D757N. The I365S + D757N ipr^HR^ haplotype (Bc-396) had a significantly lower iprodione sensitivity (EC_50_ value of 25.10 µg mL^-1^) when compared to ipr^MR^ isolates which had the I365S + V1136I haplotype (Bc-39, Bc-128, Bc- 179, Bc-343; mean EC_50_ value of 8.07 ± 1.58 µg mL^-1^) (data not shown) (Table 3 and S2, Fig. 6). Twenty-nine iprodione resistant isolates, identified in the DC assay, were tested using the CAPS method described in Oshima *et al*.^35^ (*Taq* I – I365N/R/S) and in this study (*Sma* I – Q369H/P) (Table 3). From this analysis, eighteen and eleven isolates were characterised as I365N/R/S and Q369H/P, respectively (Table 3).

Two putative pyrimethanil target genes were sequenced; *Pos5* and *Mdl1*.^16^ Four of the pyr^MR^ isolates (Bc-39, Bc-128, Bc-177, Bc-279) exhibited no changes in *Pos5* (Table S2, Fig. 6). Three other pyr^MR^ isolates (Bc-296, Bc-298, Bc-477) showed the amino acid change V273I or V273L (Tables 3 and S2, Fig. 6). All pyr^HR^ isolates had either P319A, L412F or L412V (Tables 3 and S2, Fig. 6).

With respect to the *Mdl1* gene, all isolates tested showed the amino acid changes S29P and K790R (Fig. 6). One pyr^MR^ isolate (Bc-177) and two pyr^HR^ isolates (Bc-287, Bc-289) showed the amino acid change T66A (Fig. 6). One sensitive isolate (Bc-385), one pyr^MR^ isolate (Bc-177) and two pyr^HR^ isolates (Bc-297, Bc-289) all showed the amino acid change D780N (Fig. 6).

The teb^LR^ isolate Bc-475 showed the amino acid change P347S (*K354* in the archetype *Z. tritici*) (Table 3, Fig. 6). No changes in the *Cyp51* gene were found in the other teb^LR^ isolate Bc-130. Sequencing of a 973 bp region directly upstream of the *Cyp51* start codon in Bc-130, revealed two SNPs (G > A at -365 and A > T at -169) that were not present in the sensitive comparative isolate Bc- 385 (data not shown). *Cyp51* expression analysis in Bc-130 revealed small constitutive and inducible over expression values (∼1.5-fold), compared to the comparative teb^S^ isolate Bc-385 (*P* < 0.05; Fig. S2). The amino acid changes V24I and V61I were found in the sensitive strains Bc-385 and Bc-7, respectively (Fig. 6).

### 3.4 D354Y, S611N and D616G mutations in the transcription factor gene Mrr1 may contribute to constitutive overexpression of AtrB in fludioxonil resistant isolates

The *AtrB* gene was found to be expressed constitutively at a significantly higher level in fludioxonil resistant isolates Bc-279, Bc-128 and Bc-391 compared to the flu^S^ isolates Bc-385 and Bc-287 (*P* < 0.05; Fig. 7). The relative increases in expression were 6.3, 7.3 and 9.6-fold for Bc-279, Bc-128 and Bc-391, respectively (Fig. 7). Bc-130 did not exhibit overexpression of *AtrB* compared to Bc-385 (Fig. 7). Under pyrimethanil treatment, only Bc-279, showed a significantly higher relative expression (5.9-fold) than Bc-385 (*P* < 0.05; Fig. S1). Sequencing of the MDR1-related *Mrr1* gene in the fludioxonil resistant isolates Bc-279, Bc-128, Bc-391 and the fludioxonil sensitive isolates Bc-385, Bc- 130 and Bc-287 revealed the presence of several mutations and indels (Table S5, Fig. 8). Of these mutations, V227I + S611N, D616G and Δ6bp (nt 67–72, or 73–78, or 79–84) + D354Y were only found in Bc-279, Bc-128 and Bc-391, respectively (Table S5, Fig. 8).

**Figure 7.**
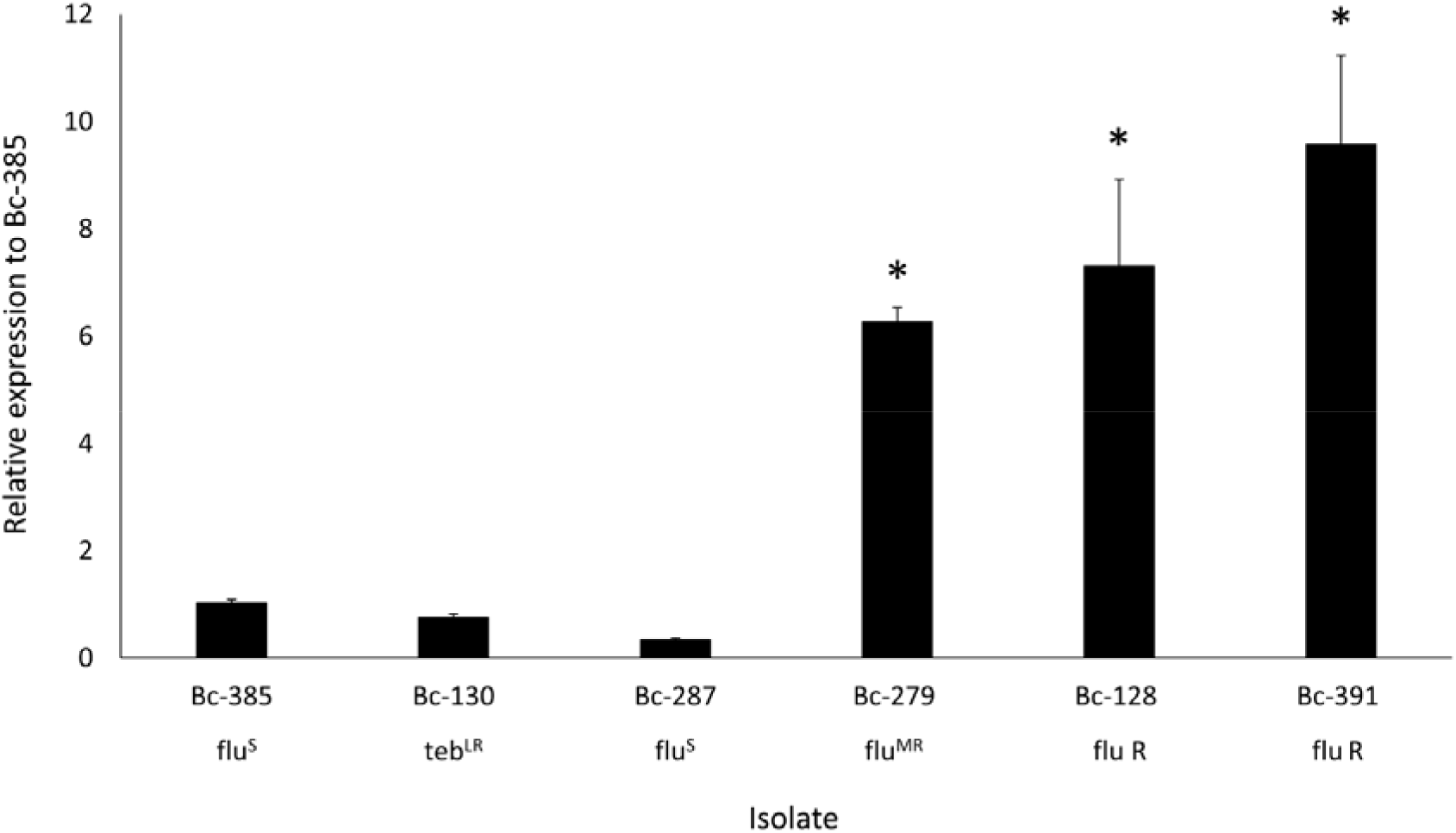
Expression analysis of *AtrB* in the sensitive comparative strain Bc-385 and resistant isolates Bc-128, Bc-130, Bc-279, Bc-287 and Bc-391. Values indicate expression levels relative to the comparative strain Bc-385 without fungicide treatment. Bc-385 untreated expression was normalised to 1. *indicates significant difference compared to Bc-385 according to an independent samples t-test (*P* = 0.05). flu = fludioxonil phenotype, teb = tebuconazole phenotype. S, LR, MR, and R indicate isolates designated as sensitive or exhibiting low resistance, medium resistance, or resistance, respectively.

## 4 Discussion

Data on fungicide resistance in Australian grapevine *B. cinerea* populations is limited.^21, 22^ To provide further data on the fungicide sensitivity status in *B. cinerea* grapevine populations, a nation- wide screening study was undertaken.

The resistance frequencies found in this study were generally low (2.1 – 11.6%) and like values previously reported for *B. cinerea* in wine grapes by other authors (Table 4, Fig. 3). Differences in frequencies among reports may be influenced by numerous contributing factors, including phenotyping methods, chemical use patterns across regions and fitness differences between *B. cinerea* strains. This study used microtiter and YSSA based mycelial growth methods to calculate overall frequencies, while previous reports used either microtiter^12, 17, 37^ or a non-YSSA based mycelial growth methods (Table 4).^13, 18, 20, 22, 38–40^

**Table 4.**
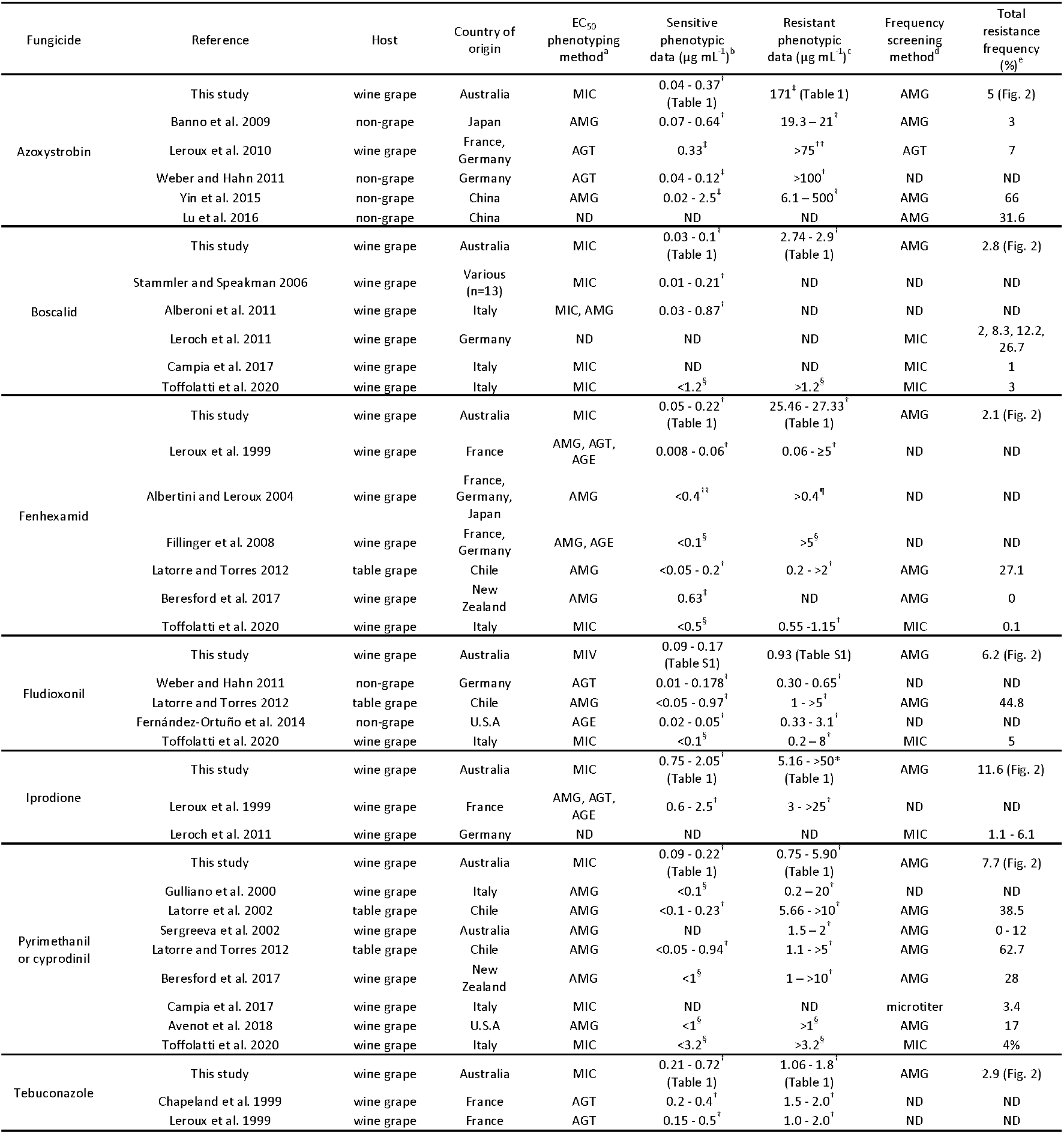
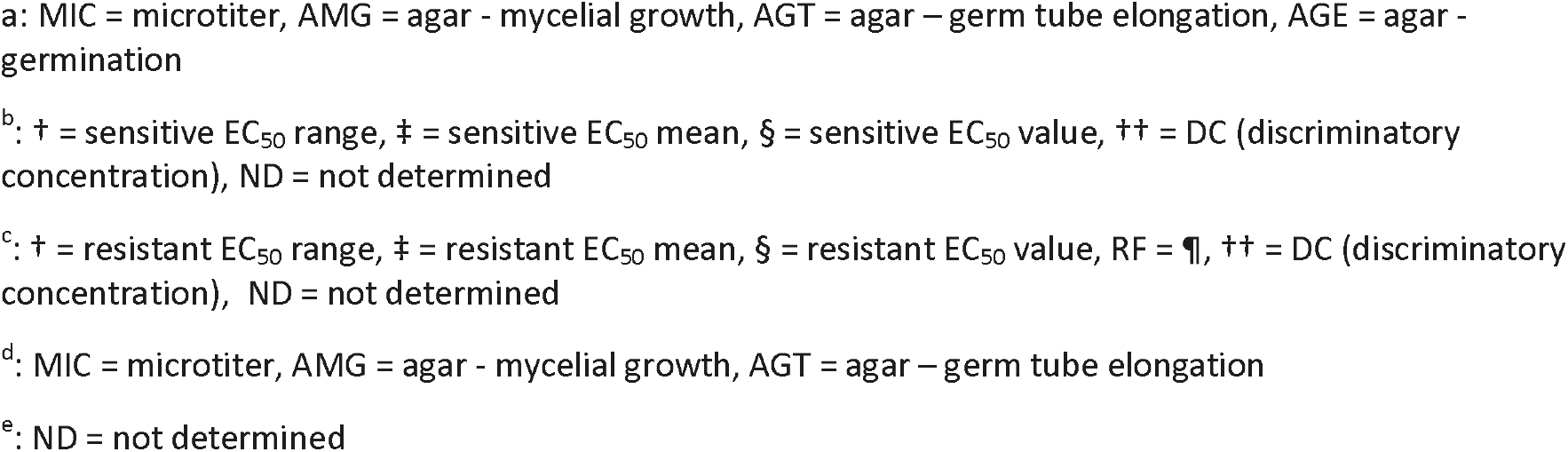
Methodologies and results for EC_50_ and frequency data from wine grape, table grape and non-grape reports for comparative purposes

Diverse chemical programs across different wine growing countries could potentially promote differences in frequencies between reports. The current recommended limit of two sprays per season per MOA in Australian wine grape production may be playing a role in limiting frequency levels across all single-site fungicides. Leroch *et al*.^12^ and Walker *et al*.^14^ characterised low iprodione resistance frequencies in *B. cinerea* and suggested that this was the consequence of limited use of these fungicides in German and French vineyards, respectively. In Australia, the relatively recent registration of fenhexamid and boscalid, together with the drastic reduction in use of the latter due to MRL restrictions, could also contribute to maintaining low resistance frequency levels for these fungicides. Conversely, the earlier introduction of pyrimethanil and iprodione could be associated with increased exposure to these chemicals and support the relatively higher resistance frequencies found. The lack of independent selection pressure for fungicides available as mixtures with other MOAs in Australia, e.g. azoxystrobin, tebuconazole and fludioxonil, could also be associated with low resistance frequency levels to these chemicals. The less frequent use of expensive, highly specific botryticides may have an effect on resistance frequencies.

Low resistance frequencies could be the result of the presence of fitness penalties. The presence of fitness costs in fenhexamid and fludioxonil resistant field isolates^27, 41–44^ could play a significant role in maintaining low resistance frequencies with respect to these fungicides. Conversely, no significant fitness costs have been reported in iprodione or pyrimethanil resistant isolates, which may support the relatively higher resistance frequency found for these two chemicals.^35, 45–49^

Resistance to multiple MOA (2 – 5) was recorded in 43.7% of the resistant isolates (Table 2). Multi-resistance has been previously reported in several wine grape *B. cinerea* studies.^13, 17, 22, 50, 51^ This phenomenon could be the result of overexposure of *Botrytis* populations to different MOAs in a sequential manner.^52^ Applications of a single MOA or mixtures of single-site actives appear to have selected for multi-resistant *B. cinerea* strains in blackberry and strawberry.^52^ Resistance to one fungicide in a multi-resistant isolate could be indirectly selected by the application of another fungicide for which resistance already exists, also known as “selection by association” theory.^53, 54^ In this report, the characterisation of multi-resistant isolates resistant to boscalid could be an example of this process as recent selection pressure from this fungicide is now essentially non-existent in *B. cinerea* populations. Generally, in Australia MRL restrictions have decreased the number of MOAs rotated within a chemical program, as alternative chemical options are absent.

Resistance identified in this study was in most cases associated with mutations reported elsewhere, with novel genotypes also identified (Fig. 6 and 8). The *CytB* G143A mutation^38^ was identified in all azo^R^ isolates (Table 3, Fig. 6). An intron found at position 143 in an azo^S^ and azo^R^ isolates (Fig. 6) had previously been described in sensitive^38, 50, 55, 56^ and QoI resistant isolates.^57^ The presence of this intron prevents the G143A mutation occurring as splicing will be affected.^38, 56, 58, 59^ In our study, the presence of this intron in an isolate (Bc-111) that also contains mutant (G143A) copies of *CytB*, confirms that Australian isolates can be heteroplasmic for the G143 intron. Intron heteroplasmy could suggest the presence of a “fitness balance”, whereby intron and G143A *CytB* copies are balanced to maintain the lowest possible fitness cost.

**Figure 8.**
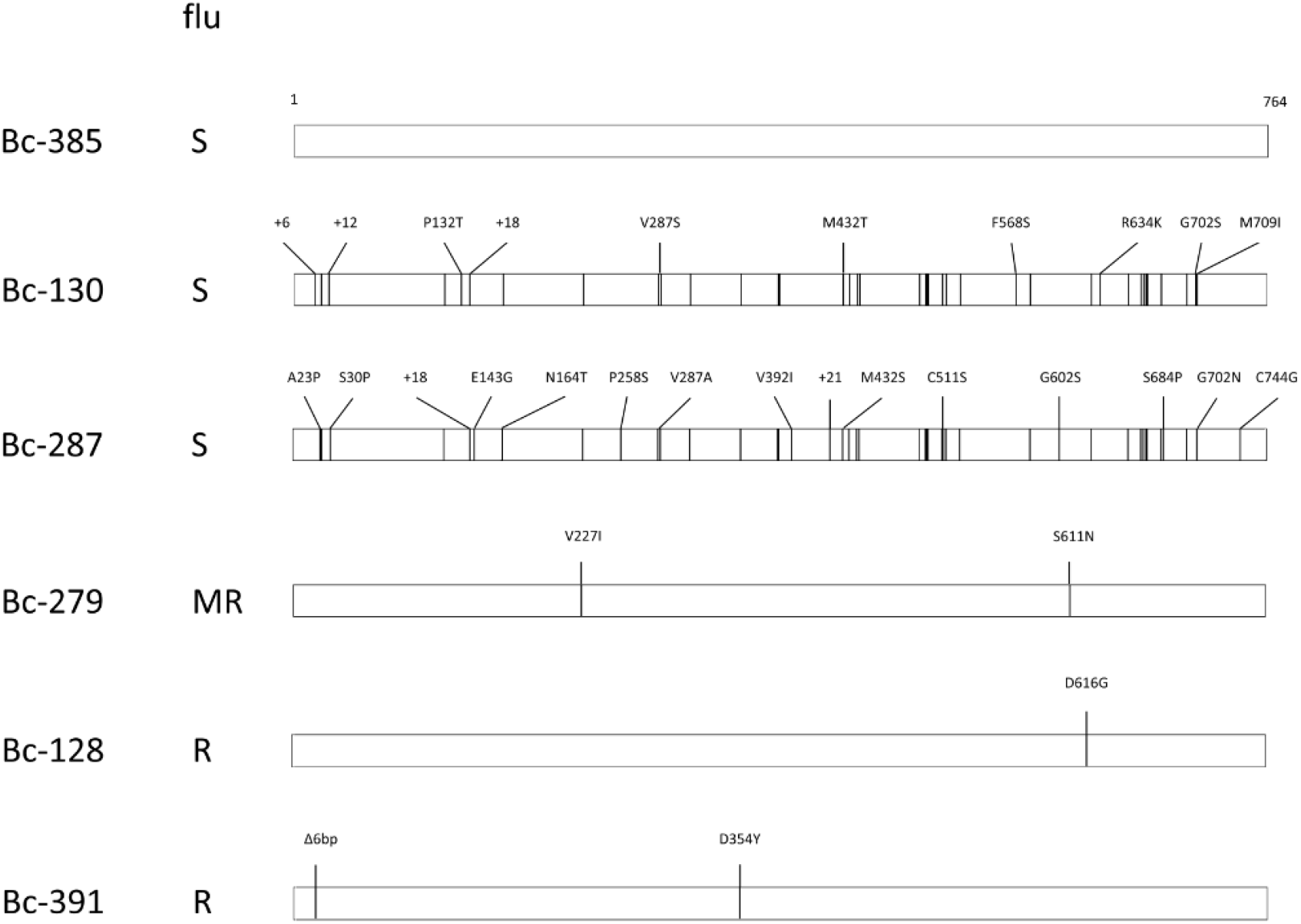
Non-synonymous changes (vertical lines) found in *Mrr1* amino acid sequences of the sensitive isolate Bc-385 and MDR1 candidate isolates; Bc-130, Bc-287, Bc-279, Bc-128 and Bc-391. All non-synonomous changes (vertical lines), insertions (+) and deletions (Δ), are indicated as compared to the reference strain B05.10. All changes unique to each isolate compared to other isolates in this study are labelled. flu = fludioxonil phenotype, S = sensitive, MR = medium resistance, R = resistance.

Mutations H272R/Y, previously found in the gene encoding for the sdhB subunit of the SDHI target in highly resistant (RF = 40) *B. cinerea*,^50, 60^ were identified in all bos^R^ isolates genotyped in this study (Table 3, Fig. 6).

Sequencing of *Erg27* in fen^HR^ isolates revealed the presence of the F412S mutation (Table 3, Fig. 6), which had been previously associated with a high level of fenhexamid resistance (RF = >30) in *B. cinerea* isolated from grapes.^31^ The P238S mutation identified in fenhexamid sensitive and resistant isolates in this study (Fig. 6), has been found in both sensitive and resistant isolates in a number of grape and non-grape studies, which suggests that P238S may not be associated with resistance.^15, 37, 61–63^

The occurrence of multi-single site resistance phenotypes in a population can disguise the presence of MDR phenotypes. Resistance to fludioxonil, has previously been shown to be associated with overexpression of the ABC transporter gene *AtrB*.^24–26, 29^ The presence of MDR1 isolates was confirmed by measuring the expression of the *AtrB* gene in isolates sensitive and resistant to fludioxonil (Fig. 7). Three different *Mrr1* haplotypes were found in MDR1 isolates (Bc-128, Bc-279 and Bc-391) overexpressing *AtrB* (Fig. 7). The *Mrr1 m*utation D354Y (Bc-391) was previously described in MDR1 isolates in two strawberry *B. cinerea* studies.^26, 52^ The 6bp deletion in Bc-391 *Mrr1* was previously reported in an MDR1 isolate alongside mutation R632I.^24^ In addition to D354Y, the novel *Mrr1* mutations S611N and D616G were found in MDR1 isolates Bc-279 and Bc-1128, respectively. Kretschmer *et al*.^29^ found the *Mrr1* mutation S611R in MDR1 isolates. It is possible that changes in the vicinity of residue 611 could have a similar effect to that of S611R. Characterisation of additional MDR1 isolates that also exhibit these changes would provide further evidence of the association of these mutations with a MDR1 phenotype. Several group S isolates, that characteristically exhibit 18 and 21 bp insertions in *Mrr1*, have previously been associated with MDR1.^24, 26, 52^ The teb^LR^ isolate Bc-130 lacked a MDR1 phenotype but exhibited a group S-like *Mrr1* haplotype, with the characteristic 18bp and 21bp INDELS present and absent, respectively (Table S5, Fig. 7 and 8).

Mosbach *et al*.^16^ associated mutations in two genes: *Pos5* and *Mdl1*, with AP resistance in grape and strawberry *B. cinerea* field isolates. All pyr^HR^ isolates showed either mutation P319A or L412F/V in *Pos5*, which agrees with the findings of Mosbach et al.,^16^ where RF values of >10 were reported in grape and strawberry *B. cinerea* isolates exhibiting these mutations.^16^ Higher EC_50_ values were found in L412V mutants than L412F mutants. (Table S2, Fig. 6). Several of the pyr^MR^ isolates exhibited V273I/L changes (Bc-296, Bc-298, Bc-477), with the remaining pyr^MR^ isolates (Bc-37, Bc- 128, Bc-177, Bc-279) showing no changes in *Pos5* (Table S2, Fig. 6). The same authors also identified AP-resistant field isolates lacking mutations in *Pos5* or *Mdl1*. Mosbach *et al*.^16^ characterised the V273I *Pos5* genotype in one AP-resistant grape isolate. Our study is the first report describing the V273L genotype, which was only found in one pyr^MR^ isolate (Fig. 6). Further screening is required to estimate the frequency of this novel genotype. A T66A change in the *Mdl1* gene was found in the pyr^MR^ in isolate Bc-177. Mosbach *et al*.^16^ characterised AP-resistant field isolates that carried T66A in combination with V273I, P319A or L412V changes in *Pos5*. The overexpression of *AtrB* in MDR1 isolates that lack *Pos5* or *Mdl1* mutations (Bc-128, Bc-279) could be contributing to the pyr^MR^ phenotype (Fig. 7). Further research is required to identify molecular markers for pyrimethanil resistance in isolates that lack mutations in *Pos5* and *Mdl1*.

The *Bos1* haplotypes I365S + V1136I (ipr^MR^), I365S + D757N (ipr^HR^) and Q369P + N373S (ipr^HR^) identified in this study have been previously associated with medium to high levels of dicarboximide resistance (RF = >3.4) in *B. cinerea* from grape and other host crops.^35, 38, 47, 64, 65^

Mutations in *Cyp51A* or *Cyp51B* are commonly associated with DMI resistance in various fungal species, however, none are yet to be identified in *B. cinerea*.^36^ A novel mutation P347S was found in the teb^LR^ isolate Bc-475, however further evidence is needed to associate this change to the teb^LR^ phenotype. The teb^LR^ isolate Bc-130 exhibited no changes in the *Cyp51* promoter or coding region (data not shown, Fig. 6). Nonetheless, expression analysis of *Cyp51* showed that Bc-130 had a marginal but significantly higher constitutive and inducible expression level of *Cyp51* when compared to the sensitive isolate Bc-385 (*P* < 0.05; Fig. S2). It is not clear whether this small difference in constitutive and inducible expression in *Cyp51* would contribute to the reduction in tebuconazole sensitivity observed in isolate Bc-130.

This study has shown that the majority of single site MOA botryticides registered in Australia for the control of botrytis bunch rot in grapes are compromised to some degree. Resistance factors are very high for some compounds and many isolates were resistant to multiple MOA across all heavily sampled Australian wine-growing regions. However, the frequency of resistance amongst isolates was low. The current recommended resistance management strategy that limits the use of single site fungicides is most likely playing a crucial role in maintaining these low frequencies.

## ACKNOWLEDGEMENTS

The authors would like to thank Steven Chang for isolating a small subset of the *B. cinerea* population, Wesley Mair for designing a selection of the oligonucleotide primers, and all growers and viticulturists who provided *Botrytis* samples. This work was funded by Wine Australia (SAR1204, SAR1701 and SAR1701-1.2) and Curtin University (Perth, Australia

## Notes

### Competing Interest Statement

The authors have declared no competing interest.

