## supplementary material for "Fungicide resistance characterised across seven modes of action in *Botrytis cinerea* isolated from Australian vineyards"

**Table S1.** Distribution of resistance across Australian wine regions.

| Wine region | State | Number of isolates tested | No. of resistant isolates^a^ | Number of vineyards sampled | Number of vineyards with resistance |
| --- | --- | --- | --- | --- | --- |
| Swan district | WA | 6 | 1 | 1 | 1 |
| Perth Hills | WA | 1 | 0 | 1 | 0 |
| Margaret River | WA | 310 | 31 | 26 | 9 |
| Geographe | WA | 2 | 2 | 1 | 1 |
| Manjimup | WA | 2 | 0 | 1 | 0 |
| Pemberton | WA | 13 | 6 | 2 | 1 |
| Great southern | WA | 30 | 16 | 7 | 5 |
| Total | WA | 364 | 56 (15.4) | 39 | 17 |
| Adelaide Hills | SA | 36 | 16 | 4 | 3 |
| Barossa Valley | SA | 4 | 0 | 1 | 0 |
| Langhorne creek | SA | 27 | 1 | 2 | 1 |
| McLaren Vale | SA | 5 | 2 | 1 | 1 |
| Riverland | SA | 3 | 0 | 1 | 0 |
| Coonawarra | SA | 19 | 4 | 3 | 3 |
| Padthaway | SA | 1 | 1 | 1 | 1 |
| Total | SA | 95 | 24 (25.3) | 13 | 9 |
| Tasmania | TAS | 21 | 13 | 1 | 1 |
| Total | TAS | 21 | 13 (61.9) | 1 | 1 |
| Beechworth | VIC | 9 | 0 | 1 | 0 |
| Gippsland | VIC | 20 | 19 | 1 | 1 |
| King Valley | VIC | 71 | 20 | 6 | 4 |
| Rutherglen | VIC | 12 | 3 | 1 | 1 |
| Yarra Valley | VIC | 10 | 3 | 1 | 1 |
| Total | VIC | 122 | 45 (36.9) | 10 | 7 |
| Hunter valley | NSW | 34 | 2 | 7 | 1 |
| Mudgee | NSW | 10 | 0 | 1 | 0 |
| Cowra | NSW | 10 | 0 | 1 | 0 |
| Orange | NSW | 22 | 1 | 1 | 1 |
| Southern Tablelands | NSW | 23 | 1 | 1 | 1 |
| Tumbarumba | NSW | 4 | 2 | 2 | 1 |
| Total | NSW | 103 | 6 (5.8) | 13 | 4 |
| Granite belt | QLD | 20 | 0 | 1 | 0 |
| Total | QLD | 20 | 0 | 1 | 0 |
| Australian total |  | 725 | 144 | 77 | 37 |

^a^Parentheses indicates percentage of resistant isolates for each state.

**Table S2.** Isolate details, EC_50_ and MIC values for the subset of 53 isolates tested within the microtiter assay and additional resistant isolates characterised in the DC agar assay.

| Isolate ID^a^ | Australian wine region, State | Year sampled | Azo  EC_50_^b^ | Azo  MIC | Boscalid  EC_50_^b^ | Bos  MIC | Fen  EC_50_^b^ | Fen  MIC | Flu  EC_50_^b^ | Flu  MIC | Ipro  EC_50_^b^ | Ipr  MIC | Pyr  EC_50_^b^ | Pyr  MIC | Teb  EC_50_^b^ | Teb  MIC |
| --- | --- | --- | --- | --- | --- | --- | --- | --- | --- | --- | --- | --- | --- | --- | --- | --- |
| Bc-7 | Manjimup, WA | 2013 | 0.07 ± 0.001 | >5 | 0.10 ± 0.003 | 1 | 0.08 ± 0.003 | 1 | 0.16 ± 0.025 | 2 | 1.55 ± 0.053 | 5 | 0.23 ± 0.012 | 1 | 0.47 ± 0.006 | 3 |
| Bc-38 | Margaret River, WA | 2013 | 0.07 ± 0.003 | 5 | 0.05 ± 0.001 | 1 | 0.07 ± 0.002 | 1 | - | - | 1.08 ± 0.057 | 3 | 0.19 ± 0.006 | 0.25 | 0.43 ± 0.019 | 3 |
| Bc-39 | Geographe, WA | 2013 | 0.31 ± 0.031 | 5 | 0.10 ± 0.005 | 1 | 0.16 ± 0.015 | >1 | - | - | **11.94 ± 0.944^MR^** | **>50** | **2.27 ± 0.165^MR^** | **10** | 0.61 ± 0.036 | 3 |
| Bc-101 | Margaret River, WA | 2013 | 0.06 ± 0.008 | 5 | 0.04 ± 0.001 | 0.1 | 0.07 ± 0.003 | 1 | - | - | 0.75 ± 0.030 | 3 | 0.14 ± 0.013 | 0.3 | 0.48 ± 0.009 | 3 |
| Bc-107 | Margaret River, WA | 2013 | 0.17 ± 0.004 | 5 | 0.07 ± 0.002 | 1 | 0.08 ± 0.005 | 1 | - | - | 1.01 ± 0.042 | 2 | 0.17 ± 0.009 | 0.4 | 0.67 ± 0.026 | 3 |
| Bc-111 | Coonawarra, SA | 2013 | **>50^R^** | **>100** | **2.74 ± 0.079^R^** | **>10** | 0.14 ± 0.006 | 1 | - | - | 1.66 ± 0.066 | 3 | 0.17 ± 0.007 | 0.3 | 0.41 ± 0.022 | 3 |
| Bc-125 | Adelaide Hills, SA | 2013 | 0.11 ± 0.031 | 5 | 0.04 ± 0.0003 | 1 | 0.05 ± 0.002 | 1 | - | - | 1.07 ± 0.083 | 2 | 0.16 ± 0.007 | 0.3 | 0.39 ± 0.020 | 3 |
| Bc-127 | Margaret River, WA | 2014 | 0.08 ± 0.014 | 5 | 0.06 ± 0.0003 | 1 | 0.09 ± 0.003 | 1 | - | - | 1.24 ± 0.069 | 3 | 0.15 ± 0.007 | 0.4 | 0.49 ± 0.013 | 3 |
| Bc-128 | Great Southern, WA | 2014 | 0.35 ± 0.019 | >5 | 0.10 ± 0.007 | 1 | 0.21 ± 0.017 | >1 | - | - | **9.34 ± 0.944^MR^** | **>50** | **0.99 ± 0.047^MR^** | **5** | 0.38 ± 0.049 | 3 |
| Bc-129 | Great Southern, WA | 2014 | 0.18 ± 0.033 | 5 | 0.04 ± 0.002 | 0.1 | 0.09 ± 0.008 | 1 | - | - | 1.06 ± 0.017 | 2 | 0.15 ± 0.017 | 0.4 | 0.40 ± 0.007 | 3 |
| Bc-130 | Great Southern, WA | 2014 | 0.37 ± 0.070 | >5 | 0.06 ± 0.003 | 1 | 0.14 ± 0.010 | 1 | - | - | 1.43 ± 0.066 | 3 | 0.10 ± 0.009 | 0.2 | **1.80 ± 0.080^LR^** | **>5** |
| Bc-144 | Great Southern, WA | 2014 | 0.09 ± 0.006 | 5 | 0.03 ± 0.003 | 0.1 | 0.07 ± 0.003 | 1 | - | - | 1.35 ± 0.045 | 3 | 0.14 ± 0.014 | 0.3 | 0.44 ± 0.032 | 3 |
| Bc-177 | Great Southern, WA | 2014 | 0.10 ± 0.013 | 5 | 0.092 ± 0.012 | 1 | 0.16 ± 0.018 | >1 | - | - | **16.78 ± 1.966^HR^** | **>50** | **0.75 ± 0.023^MR^** | **5** | 0.53 ± 0.029 | 3 |
| Bc-179 | Swan district, WA | 2014 | **>50^R^** | **>100** | 0.07 ± 0.001 | 1 | 0.10 ± 0.008 | 1 | - | - | **5.16 ± 0.231^MR^** | **25** | 0.16 ± 0.011 | 0.4 | 0.38 ± 0.026 | 3 |
| Bc-181 | Swan district, WA | 2014 | **>50^R^** | **>100** | **2.90 ± 0.169^R^** | **10** | 0.12 ± 0.005 | 1 | - | - | 1.04 ± 0.022 | 3 | 0.17 ± 0.005 | 0.6 | 0.33 ± 0.027 | 3 |
| Bc-184 | Swan district, WA | 2014 | 0.10 ± 0.006 | 5 | 0.05 ± 0.001 | 1 | 0.08 ± 0.011 | 1 | - | - | 1.19 ± 0.061 | 3 | 0.09 ± 0.007 | 0.3 | 0.44 ± 0.024 | 3 |
| Bc-197 | Pemberton, WA | 2014 | 0.04 ± 0.002 | 5 | 0.03 ± 0.002 | 0.1 | 0.07 ± 0.005 | 1 | - | - | 1.04 ± 0.068 | 3 | 0.17 ± 0.006 | 0.4 | 0.32 ± 0.018 | 3 |
| Bc-203 | Margaret River, WA | 2014 | 0.08 ± 0.003 | 5 | 0.06 ± 0.003 | 1 | 0.19 ± 0.008 | >1 | - | - | 0.97 ± 0.050 | 3 | 0.10 ± 0.012 | 0.3 | 0.48 ± 0.010 | 3 |
| Bc-215 | Granite belt, QLD | 2014 | 0.22 ± 0.022 | 5 | 0.06 ± 0.003 | 1 | 0.13 ± 0.015 | >1 | - | - | 1.21 ± 0.018 | 2 | 0.12 ± 0.003 | 0.3 | 0.44 ± 0.017 | 3 |
| Bc-232 | Perth Hills, WA | 2014 | 0.06 ± 0.003 | 5 | 0.05 ± 0.002 | 1 | 0.14 ± 0.004 | >1 | - | - | 0.99 ± 0.021 | 3 | 0.15 ± 0.018 | 0.3 | 0.27 ± 0.023 | 3 |
| Bc-247 | King valley, VIC | 2014 | ND | ND | ND | ND | ND | ND | ND | ND | ND | ND | **6.84 ± 0.571^HR^** | 25 | ND | ND |
| Bc-248 | King Valley, VIC | 2014 | 0.08 ± 0.006 | 5 | 0.05 ± 0.003 | 1 | 0.10 ± 0.003 | 1 | - | - | **28.83 ± 0.338^HR^** | **>50** | 0.11 ± 0.008 | 0.25 | 0.34 ± 0.022 | 3 |
| Bc-258 | King Valley, VIC | 2014 | 0.07 ± 0.002 | 5 | 0.06 ± 0.001 | 1 | 0.05 ± 0.006 | 1 | - | - | 1.01 ± 0.085 | 2 | 0.10 ± 0.014 | 0.4 | 0.34 ± 0.013 | 3 |
| Bc-270 | King Valley, VIC | 2014 | 0.12 ± 0.029 | 5 | 0.07 ± 0.002 | 1 | 0.08 ± 0.005 | 1 | - | - | 1.49 ± 0.023 | 2 | 0.14 ± 0.016 | 0.4 | 0.64 ± 0.124 | 3 |
| Bc-278 | King Valley, VIC | 2014 | 0.12 ± 0.014 | 5 | 0.04 ± 0.002 | 1 | 0.19 ± 0.008 | 1 | - | - | 0.96 ± 0.069 | 3 | 0.14 ± 0.009 | 0.4 | 0.40 ± 0.050 | >3 |
| Bc-279 | King Valley, VIC | 2014 | 0.23 ± 0.009 | >5 | 0.07 ± 0.003 | 1 | 0.22 ± 0.007 | >1 | **0.93 ± 0.010^MR^** | **10** | 2.05 ± 0.015 | 10 | **1.43 ± 0.104^MR^** | **5** | 0.51 ± 0.025 | 3 |
| Bc-287 | Gippsland, VIC | 2014 | 0.11 ± 0.011 | >5 | 0.05 ± 0.001 | 1 | **27.33 ± 1.879^R^** | **>100** | 0.09 ± 0.015 | 1 | **>50^HR*^HR** | **>50** | **4.66 ± 0.277^HR^** | **>10** | 0.35 ± 0.006 | 3 |
| Bc-289 | Gippsland, VIC | 2014 | 0.20 ± 0.018 | >5 | 0.05 ± 0.003 | 1 | **25.79 ± 2.531^R^** | **>100** | - | - | **40.66 ± 2.669^HR^** | **>50** | **5.90 ± 0.447^HR^** | **10** | 0.21 ± 0.014 | 3 |
| Bc-296 | Gippsland, VIC | 2014 | ND | ND | ND | ND | ND | ND | ND | ND | ND | ND | **1.09 ± 0.149^MR^** | **5** | ND | ND |
| Bc-298 | Gippsland, VIC | 2014 | ND | ND | ND | ND | ND | ND | ND | ND | ND | ND | **1.74 ± 0.142^MR^** | **5** | ND | ND |
| Bc-308 | Goulburn, NSW | 2014 | 0.08 ± 0.011 | 5 | 0.05 ± 0.003 | 1 | 0.08 ± 0.008 | 1 | - | - | 0.91 ± 0.046 | 2 | 0.12 ± 0.021 | 0.3 | 0.36 ± 0.026 | 3 |
| Bc-327 | Tasmania | 2014 | 0.09 ± 0.009 | 5 | 0.04 ± 0.001 | 1 | 0.10 ± 0.005 | 1 | - | - | 1.18 ± 0.022 | 3 | 0.14 ± 0.009 | 0.4 | 0.49 ± 0.005 | 3 |
| Bc-343 | Tasmania | 2014 | 0.09 ± 0.021 | 5 | 0.04 ± 0.001 | 1 | 0.08 ± 0.005 | 1 | - | - | **5.83 ± 0.190^MR^** | **25** | 0.15 ± 0.007 | 0.3 | 0.24 ± 0.003 | 3 |
| Bc-378 | Coonawarra, SA | 2014 | 0.07 ± 0.002 | 5 | 0.03 ± 0.002 | 0.1 | 0.05 ± 0.004 | 1 | - | - | 0.94 ± 0.028 | 3 | 0.17 ± 0.013 | 0.3 | 0.55 ± 0.026 | 3 |
| Bc-385 | McLaren Vale, SA | 2014 | 0.18 ± 0.034 | 5 | 0.06 ± 0.005 | 1 | 0.11 ± 0.007 | 1 | 0.17 ± 0.038 | 2 | 1.26 ± 0.018 | 5 | 0.15 ± 0.010 | 0.4 | 0.40 ± 0.028 | 3 |
| Bc-389 | Riverland, SA | 2014 | 0.13 ± 0.008 | 5 | 0.06 ± 0.001 | 1 | 0.09 ± 0.003 | 1 | - | - | 1.03 ± 0.042 | 3 | 0.18 ± 0.007 | 0.4 | 0.45 ± 0.016 | 3 |
| Bc-392 | Adelaide Hills, SA | 2014 | 0.13 ± 0.033 | 5 | 0.05 ± 0.003 | 0.1 | 0.09 ± 0.002 | 1 | - | - | 1.56 ± 0.047 | 2 | 0.16 ± 0.007 | 0.4 | 0.46 ± 0.031 | 3 |
| Bc-396 | Adelaide Hills, SA | 2014 | **>50^R^** | **>100** | 0.05 ± 0.002 | 1 | **25.46 ± 2.641^R^** | **>100** | - | - | **25.10 ± 0.990^HR^** | **>50** | **4.95 ± 0.560^HR^** | **>10** | 0.42 ± 0.037 | 3 |
| Bc-398 | Adelaide Hills, SA | 2014 | ND | ND | ND | ND | ND | ND | ND | ND | ND | ND | **15.21 ± 0.814^HR^** | **>25** | ND | ND |
| Bc-403 | Adelaide Hills, SA | 2014 | ND | ND | ND | ND | ND | ND | ND | ND | ND | ND | **28.71 ± 2.056^HR^** | **>25** | ND | ND |
| Bc-410 | Yarra Valley, VIC | 2015 | 0.08 ± 0.010 | 5 | 0.05 ± 0.002 | 1 | 0.10 ± 0.010 | 1 | 0.09 ± 0.007 | 1 | 1.27 ± 0.014 | 2 | 0.14 ± 0.009 | 0.3 | 0.46 ± 0.022 | 3 |
| Bc-419 | Beechworth, VIC | 2015 | 0.12 ± 0.028 | 5 | 0.05 ± 0.001 | 1 | 0.10 ± 0.007 | 1 | - | - | 1.24 ± 0.023 | 3 | 0.15 ± 0.009 | 0.3 | 0.52 ± 0.019 | 3 |
| Bc-438 | King Valley, VIC | 2015 | 0.07 ± 0.004 | 5 | 0.04 ± 0.002 | 1 | 0.06 ± 0.002 | 1 | - | - | 1.20 ± 0.042 | 2 | 0.15 ± 0.004 | 0.4 | 0.27 ± 0.023 | 3 |
| Bc-449 | Rutherglen, VIC | 2015 | 0.07 ± 0.004 | 5 | 0.04 ± 0.002 | 0.1 | 0.08 ± 0.005 | 1 | - | - | 1.17 ± 0.019 | 2 | 0.15 ± 0.018 | 1 | 0.41 ± 0.031 | 3 |
| Bc-459 | Hunter Valley, NSW | 2015 | 0.09 ± 0.009 | 5 | 0.03 ± 0.001 | 0.1 | 0.14 ± 0.003 | 1 | - | - | 1.29 ± 0.046 | 2 | 0.14 ± 0.008 | 0.25 | 0.72 ± 0.012 | 3 |
| Bc-464 | Hunter Valley, NSW | 2015 | 0.08 ± 0.005 | 5 | 0.04 ± 0.001 | 1 | 0.08 ± 0.003 | 1 | - | - | 1.04 ± 0.027 | 2 | 0.13 ± 0.002 | 0.25 | 0.50 ± 0.024 | 3 |
| Bc-472 | Hunter Valley, NSW | 2015 | 0.19 ± 0.005 | 5 | 0.04 ± 0.001 | 1 | 0.08 ± 0.002 | 1 | - | - | 1.03 ± 0.022 | 3 | 0.12 ± 0.009 | 0.25 | 0.47 ± 0.034 | 3 |
| Bc-475 | Hunter Valley, NSW | 2015 | 0.14 ± 0.028 | 5 | 0.04 ± 0.001 | 1 | 0.13 ± 0.019 | 1 | - | - | 1.40 ± 0.005 | 3 | 0.14 ± 0.010 | 0.25 | **1.06 ± 0.040^LR^** | **4** |
| Bc-477 | Hunter Valley, NSW | 2015 | ND | ND | ND | ND | ND | ND | ND | ND | ND | ND | **2.68 ± 0.174^MR^** | **10** | ND | ND |
| Bc-486 | Adelaide Hills, SA | 2015 | 0.13 ± 0.021 | 5 | 0.03 ± 0.001 | 0.1 | 0.07 ± 0.004 | 1 | - | - | 1.56 ± 0.057 | 2 | 0.14 ± 0.005 | 0.3 | 0.50 ± 0.005 | 3 |
| Bc-500 | Langhorne creek, SA | 2015 | 0.12 ± 0.029 | 5 | 0.05 ± 0.0004 | 1 | 0.12 ± 0.008 | 1 | - | - | 1.58 ± 0.047 | 2 | 0.14 ± 0.009 | 0.4 | 0.69 ± 0.029 | 3 |
| Bc-515 | Langhorne creek, SA | 2015 | 0.07 ± 0.010 | 5 | 0.05 ± 0.0005 | 1 | 0.08 ± 0.004 | 1 | - | - | 1.11 ± 0.052 | 3 | 0.12 ± 0.011 | 0.25 | 0.41 ± 0.017 | 3 |
| Bc-531 | Barossa Valley, SA | 2015 | 0.10 ± 0.018 | 5 | 0.05 ± 0.0005 | 1 | 0.09 ± 0.002 | 1 | - | - | 1.60 ± 0.040 | 3 | 0.11 ± 0.006 | 0.25 | 0.38 ± 0.046 | 3 |
| Bc-545 | Pemberton, WA | 2015 | **>50^R^** | **>100** | 0.03 ± 0.001 | 0.1 | 0.08 ± 0.005 | 1 | - | - | 1.05 ± 0.057 | 3 | 0.18 ± 0.011 | 0.4 | 0.34 ± 0.013 | 3 |
| Bc-618 | Margaret River, WA | 2015 | ND | ND | ND | ND | ND | ND | ND | ND | ND | ND | **5.09 ± 0.504^HR^** | **25** | ND | ND |
| Bc-630 | Margaret River, WA | 2015 | 0.07 ± 0.004 | 5 | 0.05 ± 0.001 | 1 | 0.07 ± 0.009 | 1 | - | - | 1.00 ± 0.013 | 2 | 0.13 ± 0.009 | 0.25 | 0.28 ± 0.017 | 3 |
| Bc-653 | Margaret River, WA | 2015 | 0.10 ± 0.006 | 5 | 0.05 ± 0.001 | 1 | 0.09 ± 0.003 | 1 | - | - | 0.80 ± 0.072 | 2 | 0.15 ± 0.004 | 0.3 | 0.28 ± 0.024 | 3 |
| Bc-655 | Great Southern, WA | 2015 | 0.10 ± 0.020 | 5 | 0.03 ± 0.002 | 0.1 | 0.12 ± 0.012 | 1 | - | - | 1.08 ± 0.049 | 3 | 0.12 ± 0.009 | 0.25 | 0.37 ± 0.031 | 3 |
| Bc-662 | Great Southern, WA | 2015 | 0.07 ± 0.003 | 5 | 0.05 ± 0.002 | 1 | 0.14 ± 0.001 | 1 | - | - | 0.76 ± 0.048 | 2 | 0.13 ± 0.019 | 0.25 | 0.41 ± 0.016 | 3 |
| Bc-665 | Adelaide Hills, SA | 2015 | 0.07 ± 0.003 | 5 | 0.04 ± 0.002 | 1 | 0.11 ± 0.007 | 1 | - | - | 1.23 ± 0.041 | 2 | 0.13 ± 0.014 | 0.25 | 0.28 ± 0.017 | 3 |

^a^: Resistant isolates Bc-247, Bc-296, Bc-298, Bc-398, Bc-403, Bc-477 and Bc-618 were characterised in the DC agar assay and subsequently phenotyped in the microtiter assay

^b^: EC_50_ values for all resistant populations were significantly difference compared to sensitive populations according to independent samples t-test (*P* = 0.05) or Mann-Whitney U-test (*P* = 0.05). LR = low resistance, MR = medium resistance, HR = high resistance, R = resistance. (-) indicates that no testing was undertaken. ND indicates that the EC_50_ was not determined

**Table S3.** List of primers, annealing temperatures and extension times used for genetic characterisation of isolates in this study

| Primer name | Primer sequence (5'-3') | Gene | Primer purpose | Annealing temperature (°C) | Extension time (min) | Reference |
| --- | --- | --- | --- | --- | --- | --- |
| cytb 139F | ACCGAATGGTGGGATCAATA | *CytB* | genotyping - flanking | 59 | 3 | This study |
| cytb 4771R | TCCGAGATACCAGTAGCGTTT | *CytB* | genotyping - flanking |  |  | This study |
| cytb 872R | ATGCCCTCAAAAGGGGATAG | *CytB* | genotyping - internal |  |  | This study |
| cytB 1017F | GGATTTACTGATGCTGAAGG | *CytB* | genotyping - internal |  |  | This study |
| cytb 1950F | AACCAACGACACACAGAAAATG | *CytB* | genotyping - internal |  |  | This study |
| cytB 1971R | CATTTTCTGTGTGTCGTTGG | *CytB* | genotyping - internal |  |  | This study |
| cytb 2408F | TCGTCGGCCATATAAAAGGT | *CytB* | genotyping - internal |  |  | This study |
| cytB 2882F | CCTAATGTTGTTACCCAAGG | *CytB* | genotyping - internal |  |  | This study |
| cytb 3154R | TTCTTCAGACATGGCTGGTC | *CytB* | genotyping - internal |  |  | This study |
| cytb 3738F | TCATTTGCAGCGAAGTTTGA | *CytB* | genotyping - internal |  |  | This study |
| cytB 4224F | CCGTGATAACAGAAATCC | *CytB* | genotyping - internal |  |  | This study |
| cytb 4832R | CCATCTCCATCCACCATACC | *CytB* | genotyping - internal |  |  | This study |
| cytB 4966R | CGAATCAAACTTCGCTGC | *CytB* | genotyping - internal |  |  | This study |
| cytB 5455R | TCTCCTAATACGTTAGGC | *CytB* | genotyping - internal |  |  | This study |
| cytB intron F | TAAAGCAACACATGGTGCCC | *CytB* | intron detection | 58 | 1 | This study |
| cytB intron R | TCAGTGTTGCCTATACCC | *CytB* | intron detection |  |  | This study |
| sdhB 1F | AGCAGTTTCGCCTTCTCC | *SdhB* | genotyping - flanking | 58 | 1.5 | This study |
| sdhB R | TAGCAATAACCGCCCAAAAC | *SdhB* | genotyping - flanking |  |  | This study |
| sdhB 1R | TAGCGTCTCTTGGAATG | *SdhB* | genotyping - internal |  |  | This study |
| sdhB F | AAGGTATCTGCGGCAGTTGTG | *SdhB* | genotyping - internal |  |  | This study |
| sdhD 1F | GCCAATCAAATCCGTTCC | *SdhD* | genotyping - flanking | 58 | 1.5 | This study |
| sdhD 1R | AATGCAATAGGGCAGAGGC | *SdhD* | genotyping - flanking |  |  | This study |
| sdhD F | CAACAAGCCCGCTGGACG | *SdhD* | genotyping - internal |  |  | This study |
| sdhD R | CTCCCCTCAAGCCCCACC | *SdhD* | genotyping - internal |  |  | This study |
| erg27 2F | GCGCTTAGGATAAATAGG | *Erg27* | genotyping - flanking | 60 | 2 | This study |
| erg27 1R | CATCACCATTATTCTGACC | *Erg27* | genotyping - flanking |  |  | This study |
| erg27 F | GGATGGGGATTGAGCGGA | *Erg27* | genotyping - internal |  |  | This study |
| erg27 int R | GAATGTTGCTCCAAGAGG | *Erg27* | genotyping - internal |  |  | This study |
| os1 2F | ATTCAAGAGCGTCATGCG | *Bos1* | genotyping - flanking | 62 | 4.5 | This study |
| os1 7R | ATGCCCATCAATCCATGC | *Bos1* | genotyping - flanking |  |  | This study |
| os1 2R | TCCTATTTCCCGTAGAGC | *Bos1* | genotyping - internal |  |  | This study |
| os1 F | TGGCCGGTGTACATGCTC | *Bos1* | genotyping – internal/CAPS | 55 | 1.5 | This study |
| os1 R | GGCGACAGCGGTGGTAAC | *Bos1* | genotyping – internal/CAPS |  |  | This study |
| os1 4F | AATTTGCGCGAGAAGTCACG | *Bos1* | genotyping - internal |  |  | This study |
| os1 4R | CCATCTGGTTGATCTTTCGC | *Bos1* | genotyping - internal |  |  | This study |
| os1 5F | CACAACGGATGTCAATACC | *Bos1* | genotyping - internal |  |  | This study |
| os1 5R | CGAAGACGGAATGAATCACC | *Bos1* | genotyping - internal |  |  | This study |
| os1 6F | ACCGTATGATCATGGAGG | *Bos1* | genotyping - internal |  |  | This study |
| os1 6R | GTGATACCAAGATCCAAAGC | *Bos1* | genotyping - internal |  |  | This study |
| os1 7F | GAACCAAGGAGAAGATTGC | *Bos1* | genotyping - internal |  |  | This study |
| cyp51 1F | CAAGTCTTCTCAACTTTCTCTTCTCT | *Cyp51* | genotyping - flanking | 58 | 2 | This study |
| cyp51 1R | TTATCGTCGCTCCCAAGCTA | *Cyp51* | genotyping - flanking |  |  | This study |
| cyp51 2F | AATGGAGGTGGAAAACTTTATGAA | *Cyp51* | genotyping - internal |  |  | This study |
| cyp51 2R | GTTTGGACAAATCCTCATACTTGAG | *Cyp51* | genotyping - internal |  |  | This study |
| cyp51 3R | CCAATGTCAGCAGTTCCCTT | *Cyp51* | genotyping - internal |  |  | This study |
| cyp51 3F | TCTACCAAGAACAAATCCAAGTCTT | *Cyp51* | genotyping - internal |  |  | This study |
| cyp51 prom F | TATCGTTGGTGGTCAGCG | *Cyp51* | genotyping - promoter | 58 | 1 | This study |
| cyp51 prom R | AGGCTCATTGGGGTTTGC | *Cyp51* | genotyping - promoter |  |  | This study |
| cyp51 q int F1 | CGATTCGATACCTCCTTTGC | *Cyp51* | RT-qPCR | 58 | 0.33 | This study |
| cyp51 q int R1 | TAGCTTGAGCACGTCGGTTT | *Cyp51* | RT-qPCR |  |  | This study |
| mrr1_TF1-1 | CCAATCATTCCCAATCATTCA | *Mrr1* | genotyping - flanking | 58 | 1 | Kretchmer *et al*.^29^ |
| mrr1_TF1-4 | GGATAGGGTATTGCGTAGATCG | *Mrr1* | genotyping - flanking |  |  | Kretchmer *et al*.^29^ |
| mrr int1F | CATCCAAGTATGACTCTCC | *Mrr1* | genotyping - internal |  |  | This study |
| mrr int1R | TTGTGCATAGTGACGACC | *Mrr1* | genotyping - internal |  |  | This study |
| mrr int2F | AAGAGCAGATGATCAAAGG | *Mrr1* | genotyping - internal |  |  | This study |
| mrr int2R | ACCAGCCATAACTTGAGCG | *Mrr1* | genotyping - internal |  |  | This study |
| mdl1 1F | CAAACATTCCAGCCGAGC | *Mdl1* | genotyping - flanking | 62 | 3.5 | This study |
| mdl1 6R | CAAAGTATACTGGCGAGC | *Mdl1* | genotyping - flanking |  |  | This study |
| mdl1 2F | GGAGAACGAATTGTAGCC | *Mdl1* | genotyping - internal |  |  | This study |
| mdl1 3R | GATCTCCAACTCGATTGG | *Mdl1* | genotyping - internal |  |  | This study |
| mdl1 4F | AAACCCCATGCATCAAGG | *Mdl1* | genotyping - internal |  |  | This study |
| mdl1 5R | TGTATCCAGACCTTCAGG | *Mdl1* | genotyping - internal |  |  | This study |
| pos5 1F | TTGCTAGATCGAGCCTTGC | *Pos5* | genotyping - flanking | 57 | 3 | This study |
| pos5 6R | GTCACGTTGTTCCATTTGG | *Pos5* | genotyping - flanking |  |  | This study |
| pos5 2F | GTACTCTAGGGTTCTTGG | *Pos5* | genotyping - internal |  |  | This study |
| pos5 3R | GTTTGGATGTGAGGATGC | *Pos5* | genotyping - internal |  |  | This study |
| pos5 4F | CAAGAGGTATGTTTCCGC | *Pos5* | genotyping - internal |  |  | This study |
| pos5 5R | AAATAGCAGGGTTTCTGGG | *Pos5* | genotyping - internal |  |  | This study |
| atrBfor | GCACTTGTGGCGAGTATCTATC | *AtrB* | RT-qPCR | 58 | 0.33 | Kretchmer *et al*.^29^ |
| atrBrev | TGCATCCCTCCATCCATAGC | *AtrB* | RT-qPCR |  |  | Kretchmer *et al*.^29^ |
| actin for | TCTGTCTTGGGTCTTGAGAG | *Actin* | RT-qPCR | 58 | 0.33 | Kretchmer *et al*.^29^ |
| actin rev | GGTGCAAGAGCAGTGATTTC | *Actin* | RT-qPCR |  |  | Kretchmer *et al*.^29^ |

**Table S4.** Genbank accession numbers for all DNA sequences produced from genotyping of comparative sensitive and resistant isolates in this study.

| Isolate ID | Australian wine region, State | Year sampled | *cytB^a^* | *sdhB^a^* | *sdhD^a^* | *erg27^a^* | *mrr1^a^* | *bos1^a^* | *mdl1^a^* | *pos5^a^* | *cyp51^a^* |
| --- | --- | --- | --- | --- | --- | --- | --- | --- | --- | --- | --- |
| Bc-7 | Manjimup, WA | 2013 | MW090081 | MW090065 | MW090076 | MW090088 | ND | MW090099 | MW090111 | MW090121 | MW114677 |
| Bc-39 | Geographe, WA | 2013 | ND | ND | ND | ND | ND | MW090100 | MW090112 | MW090122 | ND |
| Bc-111 | Coonawarra, SA | 2013 | ND | MW090066 | MW090077 | ND | ND | ND | ND | ND | ND |
| Bc-128 | Great Southern, WA | 2014 | ND | ND | ND | ND | MW090094 | MW090101 | MW090113 | MW090123 | ND |
| Bc-129 | Great Southern, WA | 2014 | ND | ND | ND | ND | ND | ND | ND | ND | ND |
| Bc-130 | Great Southern, WA | 2014 | ND | ND | ND | ND | ND | ND | ND | ND | MW114678 |
| Bc-175 | Great southern, WA | 2014 | ND | MW090067 | ND | ND | ND | ND | ND | ND | ND |
| Bc-177 | Great Southern, WA | 2014 | ND | ND | ND | ND | ND | MW090102 | MW090114 | MW090124 | ND |
| Bc-179 | Swan district, WA | 2014 | MW090082 | ND | ND | ND | ND | MW090103 | ND | ND | ND |
| Bc-181 | Swan district, WA | 2014 | MW090083 | MW090068 | MW090078 | ND | ND | ND | ND | ND | ND |
| Bc-247 | Beechworth, VIC | 2014 | ND | MW09069 | ND | ND | ND | ND | ND | MZ326713 | ND |
| Bc-248 | King Valley, VIC | 2014 | ND | ND | ND | ND | ND | MW090104 | ND | ND | ND |
| Bc-279 | King Valley, VIC | 2014 | ND | ND | ND | ND | MW090095 | ND | MW090115 | MW090125 | ND |
| Bc-287 | Gippsland, VIC | 2014 | ND | ND | ND | MW090089 | MW090096 | MW090105 | MW090116 | MW090126 | ND |
| Bc-289 | Gippsland, VIC | 2014 | ND | ND | ND | MW090090 | ND | MW090106 | MW090117 | MW090127 | ND |
| Bc-296 | Gippsland, VIC | 2014 | ND | ND | ND | ND | ND | ND | ND | MZ326714 | ND |
| Bc-298 | Gippsland, VIC | 2014 | ND | ND | ND | ND | ND | ND | ND | MZ326715 | ND |
| Bc-343 | Tasmania | 2014 | ND | ND | ND | ND | ND | MW090107 | ND | ND | ND |
| Bc-385 | McLaren Vale, SA | 2014 | MW090087 | MW090070 | MW090079 | MW090091 | MW090097 | MW090108 | MW090118 | MW090128 | MW114679 |
| Bc-388 | McLaren Vale, SA | 2014 | ND | MW090071 | ND | ND | ND | ND | ND | ND | ND |
| Bc-391 | Adelaide Hills, SA | 2014 | ND | ND | ND | ND | MW090098 | ND | ND | ND | ND |
| Bc-396 | Adelaide Hills, SA | 2014 | MW090084 | ND | ND | MW090092 | ND | MW090109 | MW090119 | MW090129 | ND |
| Bc-398 | Adelaide Hills, SA | 2014 | ND | ND | ND | ND | ND | ND | ND | MZ326716 | ND |
| Bc-403 | Adelaide Hills, SA | 2014 | ND | MW090072 | ND | ND | ND | ND | ND | MZ326717 | ND |
| Bc-410 | Yarra Valley, VIC | 2015 | MW090085 | MW090073 | MW090080 | MW090093 | ND | MW090110 | MW090120 | MW090130 | MW114680 |
| Bc-475 | Hunter Valley, NSW | 2015 | ND | ND | ND | ND | ND | ND | ND | ND | MW114681 |
| Bc-477 | Hunter Valley, NSW | 2015 | ND | ND | ND | ND | ND | ND | ND | MZ326718 | ND |
| Bc-545 | Pemberton, WA | 2015 | MW090086 | ND | ND | ND | ND | ND | ND | ND | ND |
| Bc-612 | Margaret River, WA | 2015 | ND | MW090074 | ND | ND | ND | ND | ND | ND | ND |
| Bc-614 | Margaret River, WA | 2015 | ND | MW090075 | ND | ND | ND | ND | ND | ND | ND |
| Bc-618 | Margaret River, WA | 2015 | ND | ND | ND | ND | ND | ND | ND | MZ326719 | ND |

^a^ND indicates that the sequence was not determined

**Table S5.** All changes found in the *mrr1* gene for sensitive and comparative MDR1 candidate isolates.

| Non-synonymous change/insertion/deletions | Nucleotide change | Bc-385  flu^S a^ | Bc-130  teb^LR a^ | Bc-287  flu^S a^ | Bc-279  flu^MR a^ | Bc-128  flu R^a^ | Bc-391  flu R^a^ |
| --- | --- | --- | --- | --- | --- | --- | --- |
| ::6bp | Between nt 51 - 52 | - | + | - | - | - | - |
| Δ6bp | nt 67 – 72, or 73 – 78, or 79 - 84 | - | - | - | - | - | + |
| A22P | GCA to CCG | - | + | + | - | - | - |
| A23P | GCT to CCA | - | - | + | - | - | - |
| ::12bp | Between nt 84 - 85 | - | + | - | - | - | - |
| S30P | TCA to CCA | - | - | + | - | - | - |
| Y119D | TAC to GAC | - | + | + | - | - | - |
| P132T | CCT to ACT | - | + | - | - | - | - |
| ::18bp | nt494 | - | + | + | - | - | - |
| E143G | GAG to GGG | - | - | + | - | - | - |
| N164T | AAC to ACT | - | - | + | - | - | - |
| V227I | GTC to ATC | - | - | - | + | - | - |
| I228T | ATA to ACA | - | + | + | - | - | - |
| P258S | CCA to TCA | - | - | + | - | - | - |
| V287A/S | GTG to GCA/TCA | - | V287S | V287A | - | - | - |
| A289S | GCT to TCT | - | + | + | - | - | - |
| N312Q | AAT to CAG | - | + | + | - | - | - |
| D354Y | GAT to TAT | - | - | - | - | - | + |
| T352A | ACT to GCT | - | + | + | - | - | - |
| Q381K | CAG to AAG | - | + | + | - | - | - |
| Q382E | CAA to GAA | - | + | + | - | - | - |
| V392I | GTG to ATA | - | - | + | - | - | - |
| ::21bp | Nt1268 | - | - | + | - | - | - |
| M432S/T | ATG to TCG/ACG | - | M432T | M432S | - | - | - |
| S437T | AGT to ACT | - | + | + | - | - | - |
| I443L | ATA to TTA | - | + | + | - | - | - |
| I445F | ATC to TTC | - | + | + | - | - | - |
| I492V | ATC to GTT/GTC | - | + | + | - | - | - |
| L497V | CTC to GTC | - | + | + | - | - | - |
| A498T | GCA to ACA | - | + | + | - | - | - |
| G499C | GGT to TGT | - | + | + | - | - | - |
| Y510F | TAT to TTT | - | + | + | - | - | - |
| C511S | TGT to TCT | - | - | + | - | - | - |
| I513V | ATA to GTA | - | + | + | - | - | - |
| V524A | GTG to GCG | - | + | + | - | - | - |
| F568S | TTC to TCC | - | + | - | - | - | - |
| V579A | GTA to GCA | - | + | + | - | - | - |
| E601G | GGC to GGG | - | + | + | - | - | - |
| G602S | GGC to AGC | - | - | + | - | - | - |
| S611N | AGT to AAT | - | - | - | + | - | - |
| D616G | GAT to GGT | - | - | - | - | + | - |
| R627K | AGG to AAG | - | + | + | - | - | - |
| R634K | AGA to AAA | - | + | - | - | - | - |
| R656L | CGC to CTC | - | + | + | - | - | - |
| N666D | AAT to GAT | - | + | + | - | - | - |
| A668G | GCT to GGT | - | + | + | - | - | - |
| G670E | GCT to GGT | - | + | + | - | - | - |
| C671F | TGC to TTC | - | + | + | - | - | - |
| C682R | TGT to CGT | - | + | + | - | - | - |
| S684P | TCT to CCT | - | - | + | - | - | - |
| G702N/S | GGC to AAC/AGC | - | G702S | G702N | - | - | - |
| M709I | ATG to ATA | - | + | - | - | - | - |
| G710C | GGT to TGT | - | + | + | - | - | - |
| C744G | TGT to GGT | - | - | + | - | - | - |
| Y760H | TAT to CAT | - | - | - | - | - | - |

^a^: + or – indicates presence or lack of a particular change. All changes are relative to the sensitive reference strain B05.10. ^S^ = sensitive, ^LR^ = low resistance, ^MR^ = high resistance, R = resistance

**Supplementary figure legends**

**Figure S1.** Expression analysis of *AtrB* in the sensitive comparative strain Bc-385 and resistant isolates Bc-287 (flu^S^ pyr^HR^) and Bc-279 (flu^MR^ pyr^MR^). flu = fludioxinil treatment, pyr = pyrimethanil treatment. Values indicate expression levels relative to the comparative strain Bc-385 without fungicide treatment. Bc-385 untreated expression was normalised to 1. *indicates significant difference compared to Bc-385 pyrimethanil treated according to an independent samples t-test (*P* = 0.05).

**Figure S2.** Expression analysis of *Cyp51* in the sensitive comparative strain Bc-385 and teb^LR^ isolate Bc-130. teb = tebuconazole treatment. Values indicate expression levels relative to the comparative strain Bc-385 untreated. Bc-385 untreated expression was normalised to 1. *indicates significant difference compared to respective Bc-385 treatment according to an independent samples t-test (*P* = 0.05).

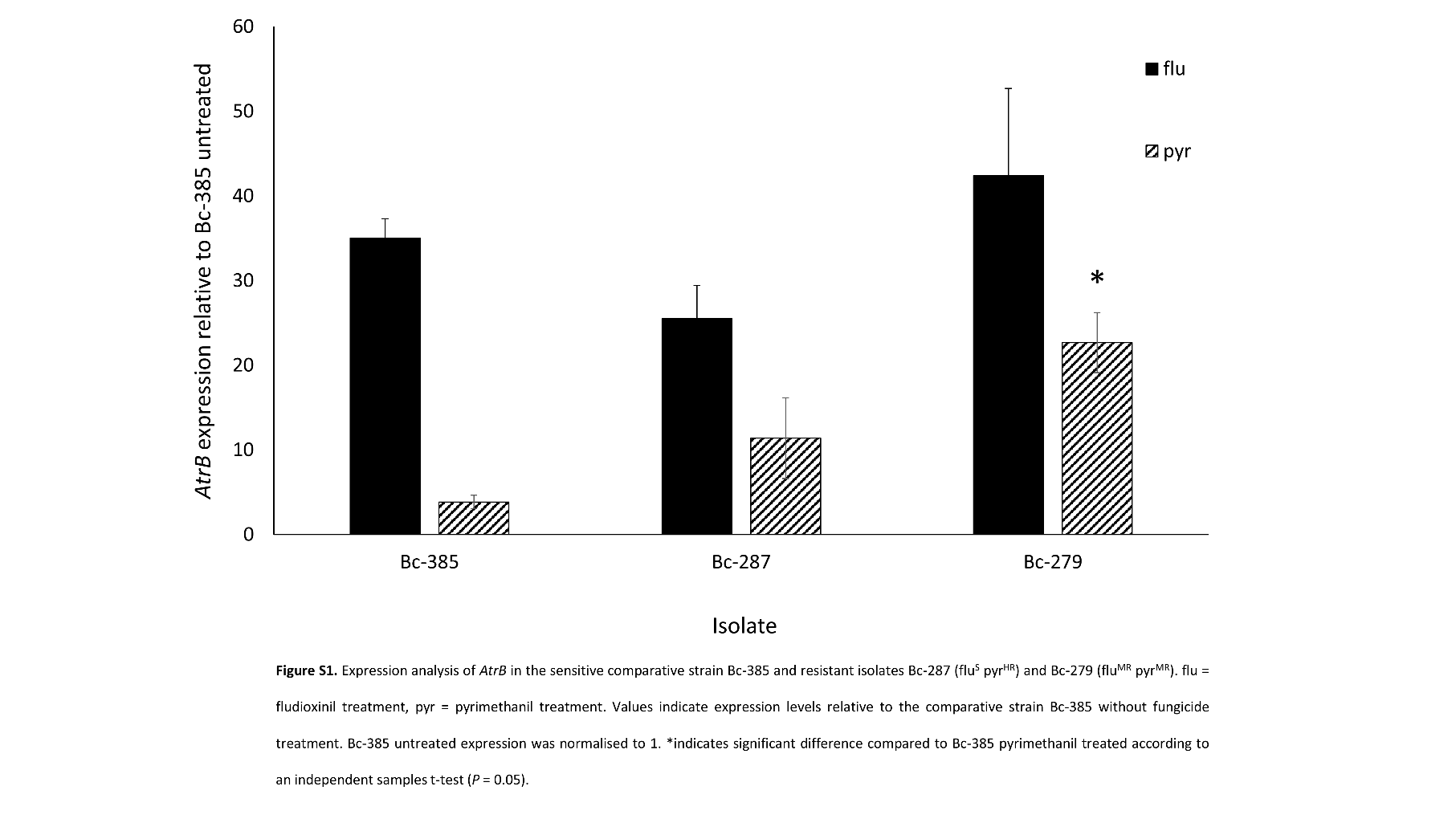

**Figure S1.** Expression analysis of *AtrB* in the sensitive comparative strain Bc-385 and resistant isolates Bc-287 (flu^S^ pyr^HR^) and Bc-279 (flu^MR^ pyr^MR^). flu = fludioxinil treatment, pyr = pyrimethanil treatment. Values indicate expression levels relative to the comparative strain Bc-385 without fungicide treatment. Bc-385 untreated expression was normalised to 1. *indicates significant difference compared to Bc-385 pyrimethanil treated according to an independent samples t-test (*P* = 0.05).

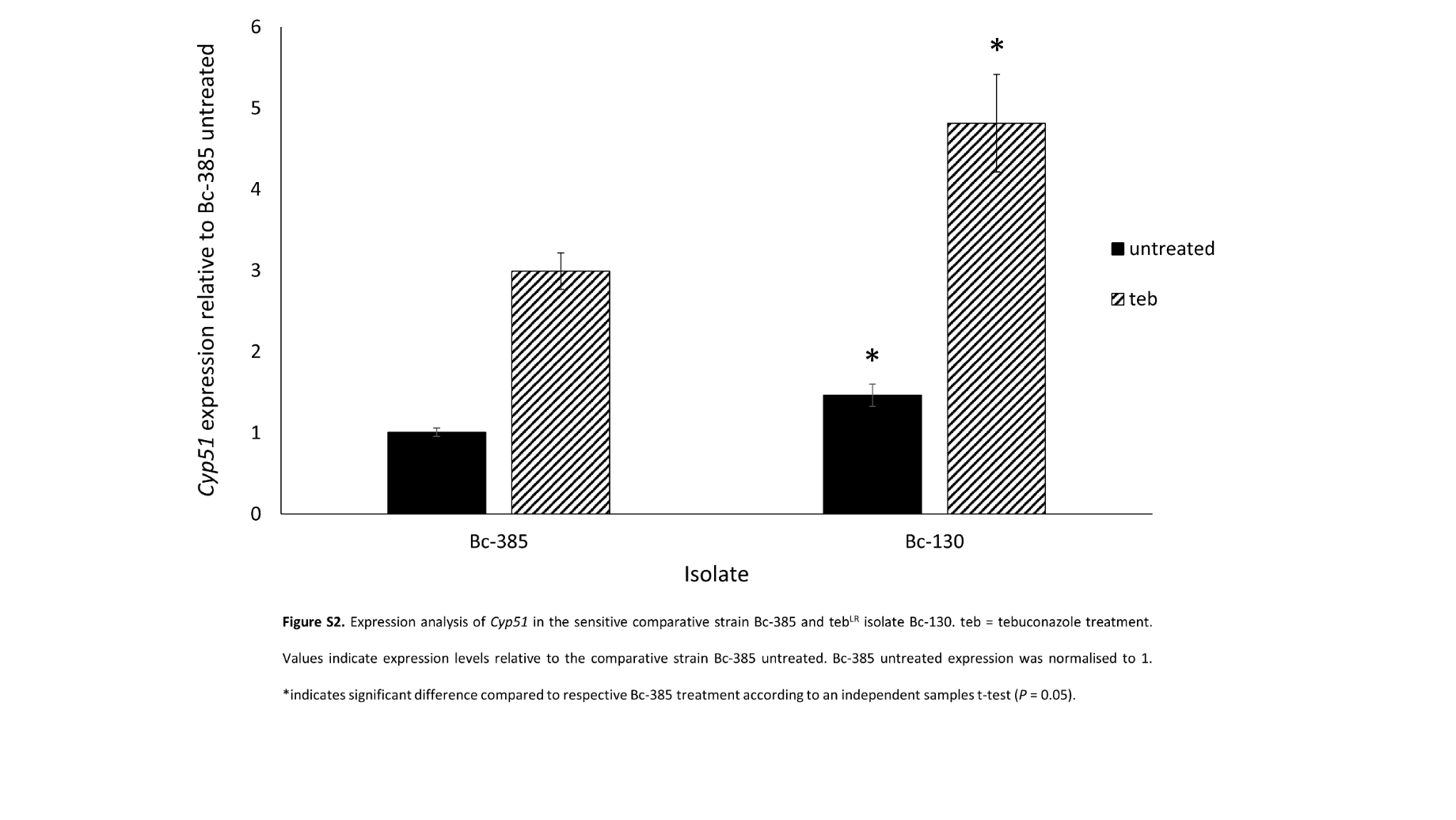

**Figure S2.** Expression analysis of *Cyp51* in the sensitive comparative strain Bc-385 and teb^LR^ isolate Bc-130. teb = tebuconazole treatment. Values indicate expression levels relative to the comparative strain Bc-385 untreated. Bc-385 untreated expression was normalised to 1. *indicates significant difference compared to respective Bc-385 treatment according to an independent samples t-test (*P* = 0.05).
